# Profiling transcription factor sub-networks in type I interferon signaling and in response to SARS-CoV-2 infection

**DOI:** 10.1101/2021.01.25.428122

**Authors:** Chilakamarti V. Ramana

**Affiliations:** Dartmouth-Hitchcock Medical Center, Dartmouth Med School, Lebanon, New Hampshire

**Keywords:** Interferon-α/β, Jak-Stat, Transcription factor, Protein-protein interaction, Signal transduction pathway, Coronaviruses, COVID-19

## Abstract

Type I interferons (IFN α/β) play a central role in innate immunity to respiratory viruses, including coronaviruses. Genetic defects in type I interferon signaling were reported in a significant proportion of critically ill COVID-19 patients. Extensive studies on interferon-induced intracellular signal transduction pathways led to the elucidation of the Jak-Stat pathway. Furthermore, advances in gene expression profiling by microarrays have revealed that type I interferon rapidly induced multiple transcription factor mRNA levels. In this study, transcription factor profiling in the transcriptome was used to gain novel insights into the role of inducible transcription factors in response to type I interferon signaling in immune cells and in lung epithelial cells after SARS-CoV-2 infection. Modeling the interferon-inducible transcription factor mRNA data in terms of distinct sub-networks based on biological functions such as antiviral response, immune modulation, and cell growth revealed enrichment of specific transcription factors in mouse and human immune cells. The evolutionarily conserved core type I interferon gene expression consists of the inducible transcriptional factor mRNA of the antiviral response sub-network and enriched in granulocytes. Analysis of the type I interferon-inducible transcription factor sub-networks as distinct protein-protein interaction pathways revealed insights into the role of critical hubs in signaling. Interrogation of multiple microarray datasets revealed that SARS-CoV-2 induced high levels of IFN-beta and interferon-inducible transcription factor mRNA in human lung epithelial cells. Transcription factor mRNA of the three major sub-networks regulating antiviral, immune modulation, and cell growth were differentially regulated in human lung epithelial cell lines after SARS-CoV-2 infection and in the tissue samples of COVID-19 patients. A subset of type I interferon-inducible transcription factors and inflammatory mediators were specifically enriched in the lungs and neutrophils of COVID-19 patients. The emerging complex picture of type I IFN transcriptional regulation consists of a rapid transcriptional switch mediated by the Jak-Stat cascade and a graded output of the inducible transcription factor activation that enables temporal regulation of gene expression.

## 1 Introduction

Interferons (IFN) are pleiotropic cytokines and exert a wide range of biological activities that include antiviral, antiproliferative, and immunoregulatory effects (1-3). There are 2 major classes of type I interferons consisting of IFN-alpha (IFN-α) represented by 14 isoforms and one IFN-beta (IFN-β). Type I IFN is produced by many cell types, including leukocytes, dendritic cells, and fibroblasts. The amount of IFN-α vs IFN-β produced varies depending on cell type and also on the virus/stimulus (1-4). The biological effects of interferons are mainly mediated by the rapid and dramatic changes in gene expression of several hundred genes (5). The role of the Janus kinase and signal transducers and activators of transcription (Jak-Stat) pathway and transcriptional regulation by interferon-stimulated gene complex (ISGF-3) consisting of Stat1, Stat2 and, Irf-9 in Type I IFN signaling has been well established (1-6). Recent advances in high throughput gene expression profiling in primary immune cells have shown that IFN α/β induces several transcription factor mRNA that sustains secondary and tertiary rounds of transcription (7,8). Several studies had shown that the activation of the canonical Jak-Stat pathway is not sufficient to explain the broad range of biological activities of type I IFN signaling. IFN α/β inhibits the interleukin-7-mediated growth and survival of B lymphoid progenitors by a Stat1-independent pathway (9). This growth inhibitory effect involves cell cycle arrest followed by apoptosis and mediated by Stat1-independent induction of Daxx, a Fas binding protein implicated in apoptosis (10). The most prominent phenotype of the Stat1-deficient mice is their increased susceptibility to microbial and viral infection due to the decreased ability to respond to the antiviral effects of the interferons (11,12). Nevertheless, Stat1-deficient mice mount an IFN-mediated resistance to virus infection. Stat1-null mice are more resistant to challenge with murine cytomegalovirus (MCMV) or Sindbis virus than mice lacking both the type I and type II IFN receptors (13). There is evidence for differential regulation of type I interferon signaling and interferon regulatory factors in neuronal cells and astrocytes of wild-type and Stat1-deficient mice in the mouse brain (14,15). IFN-α/β regulates dendritic cells (DC) and natural killer (NK) cells that are resident immune cells of the lung and are critical for innate immunity to respiratory viruses (16,17). The availability of gene expression datasets of primary immune cells treated with Interferon α/β for a short period of time facilitated profiling type I interferon-inducible transcription factor mRNA in the transcriptome (7.8). Coronaviruses are RNA viruses of the Coronaviridae family, including Severe Acute Respiratory Syndrome coronaviruses SARS-CoV-1 and SARS CoV-2 (18). The current pandemic of coronavirus disease known as COVID-19 is caused by a highly infectious SARS-CoV-2 that emerged first in Wuhan, China (19,20). The world health organization (WHO) has declared the current pandemic as a global public health emergency because of the rapid spread around the world with high mortality and morbidity. The importance of a functional interferon system for protection against SARS-CoV-2 was demonstrated by the report that 10% of nearly a thousand Covid-19 patients who developed fatal pneumonia had autoantibodies to interferons and an additional 3-5% of critically ill patients had mutations in genes that control interferons (21,22). In addition to the genetic deficiency, genetic variants in the components of type I Interferon signaling such as type I IFN receptor (IFNAR2), protein tyrosine kinase (TYK2) and the antiviral target gene Oligoadenylate synthetase (OAS1) were detected in critically ill COVID19 patients (23). Transcription factor profiling is a powerful technique to gain insights into mammalian signal transduction pathways. (24,25). Transcription factor profiling of genes involved in a biologic function such as innate and adaptive immunity led to the identification of functional connectivity of signaling hubs, critical nodes, and modules (25,26). In this study transcription factor profiling in the transcriptome of immune cells in response to type I Interferon treatment and lung epithelial cells in response to SARS-CoV-2 infection was investigated.

## 2 Materials and Methods

### Gene expression datasets

Gene expression profiling in response to Type I interferon treatment in human peripheral blood mononuclear cells (PBMC) and mouse immune cells were published previously (7,8). Gene expression datasets representing human lung cell lines infected with coronaviruses and tissue samples of healthy and COVID-19 patients were also reported (27,28). Supplementary data was downloaded from the Journal publisher websites and from Geo datasets that were archived at Pubmed (NCBI). The identification of the differentially expressed genes in the transcription profile was analyzed using the GEO2R tool and differential expression analysis using DESeq2 using default parameters. Gene expression resources from Immgen RNA seq SKYLINE COVID-19 resources were used (http://rstats.immgen.org/Skyline_COVID-19/skyline.html). Cluster analysis was performed using gene expression software tools (29). Outliers of expression were not included in the analysis. The evaluation of the gene ontology (GO) enrichment and signaling pathway analysis was conducted using DAVID, KEGG, and Metascape software (30-32). Protein interactions were interrogated and visualized in the STRING, WIKI-PI, and BIOGRID databases (33-35) Protein-protein interaction network centrality features were calculated according to https://www.sscnet.ucla.edu/soc/faculty/mcfarland/soc112/cent-ans.htm

## 3 Results and Discussion

### 3.1 Functional Annotation of the Inducible transcription factors of Type I Interferon signaling

Gene expression profiling studies in human peripheral blood monocytic cells (PBMC) and mouse immune cells revealed that a large number of transcription factor mRNA were significantly induced in the transcriptome by type I interferons (7,8). A master list of 35 human and 32 mouse interferon-induced transcription factors was subjected to the functional annotation in Database for Annotation, Visualization and Integrated Discovery (DAVID), Kyoto Encyclopedia of Genes and Genomes (KEGG), and Metascape to investigate underlying biological processes and signaling pathways (30-32). These transcription factors were broadly organized into three major functional categories-antiviral response, immune modulation, and cell growth (Table1). Mapping the list of interferon-induced transcription factors into gene ontogeny (GO) terms in Metascape provided insights into the common functional categories and signal transduction pathways in the human and mouse transcriptomes (Figure 1A and 1B). These GO terms included human papillomavirus infection, Hepatitis B, HTLV-1, response to the virus, interferon-gamma mediated signaling pathway in the antiviral response category. Immune modulation or inflammation category GO terms included regulation of cytokine production, Toll-like receptor signaling pathway, and AP1 pathway. In contrast, cell growth terms included transcriptional mis-regulation in cancer, positive regulation of cell death, and DNA damage response (Figure 1A and 1B). Protein-protein interaction abundance analysis using molecular complex detection (MCODE) algorithm in Metascape revealed three modules consisting of cell proliferation/Ionizing radiation, interferon-γ and interferon-α/β signaling represented by red, blue and green, respectively (Figure 1C). The human type I interferon-induced transcription factor data was obtained from gene expression profiling of a heterogeneous mixture of cells from the PBMC while the mouse list was derived from a purified population of B-lymphocytes, dendritic cells, granulocytes, natural killer cells, macrophages, and T-lymphocytes. Cluster analysis in gene expression profiling is often used to discriminate genes that are co-regulated under the given experimental conditions (29). The grouping of transcription factors into functional sub-categories facilitated the identification of enrichment in distinct cell types. For example, mouse antiviral transcription factors were highly expressed in granulocytes among immune cells (Figure 2A). Granulocytes including neutrophils, eosinophils, and basophils are characterized by the presence of large cytoplasmic granules and essential for the control of infection. Enhanced expression of transcription factors involved in inflammation such as Ahr, Bcl3. and Egr2 in NK cells and cell growth such as Myc, Max, and Jun in B-cells was observed (Figure 2B and Figure 2C). Furthermore, IFN-α and IFN-β mediated changes in distinct functional sub-categories such as antiviral response, immune modulation and cell growth of human PBMC can be compared in detail (Figure 3).

**Table 1.**
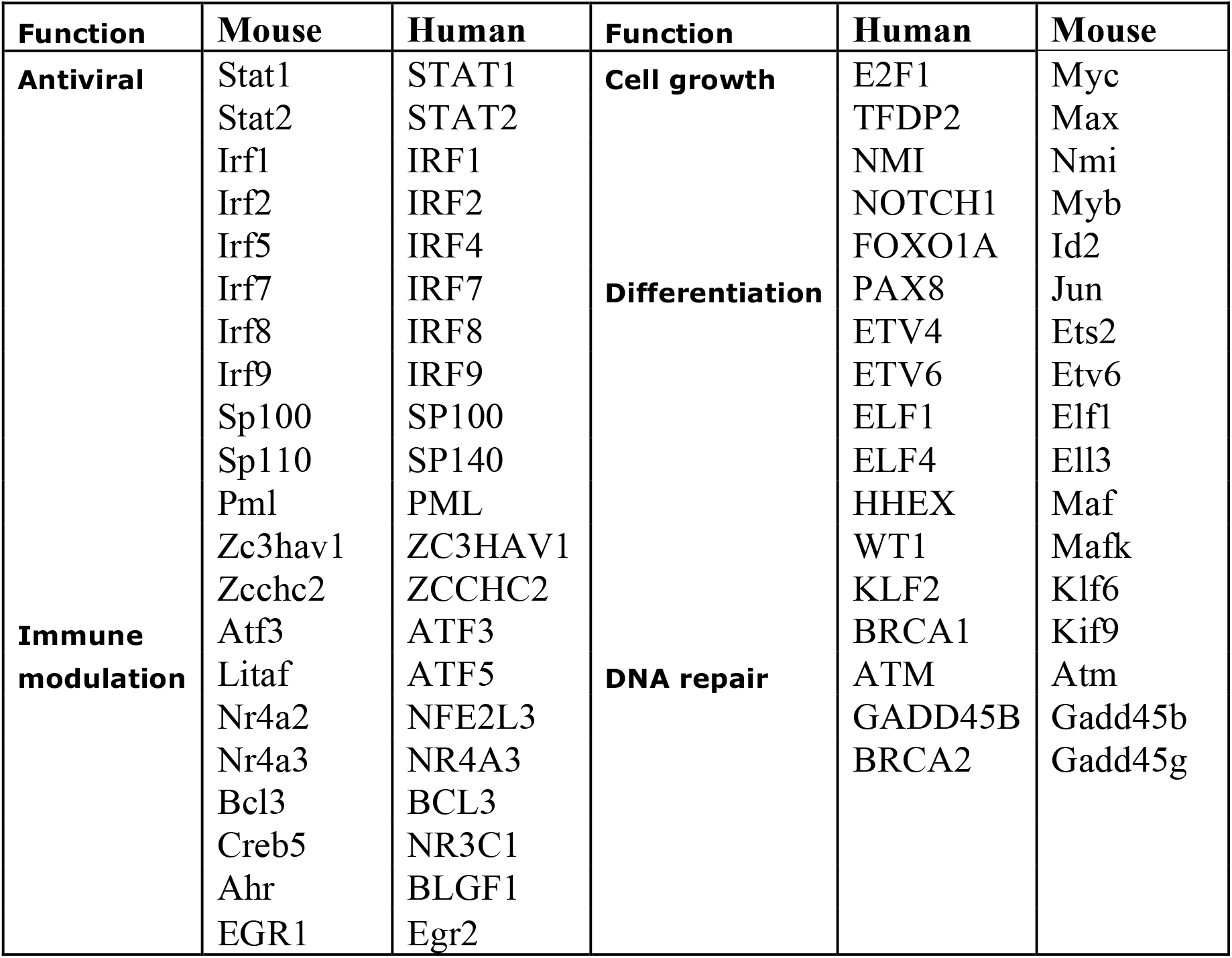
Inducible transcription factors in Type I Interferon signaling in human and mouse immune cells

**Figure 1.**
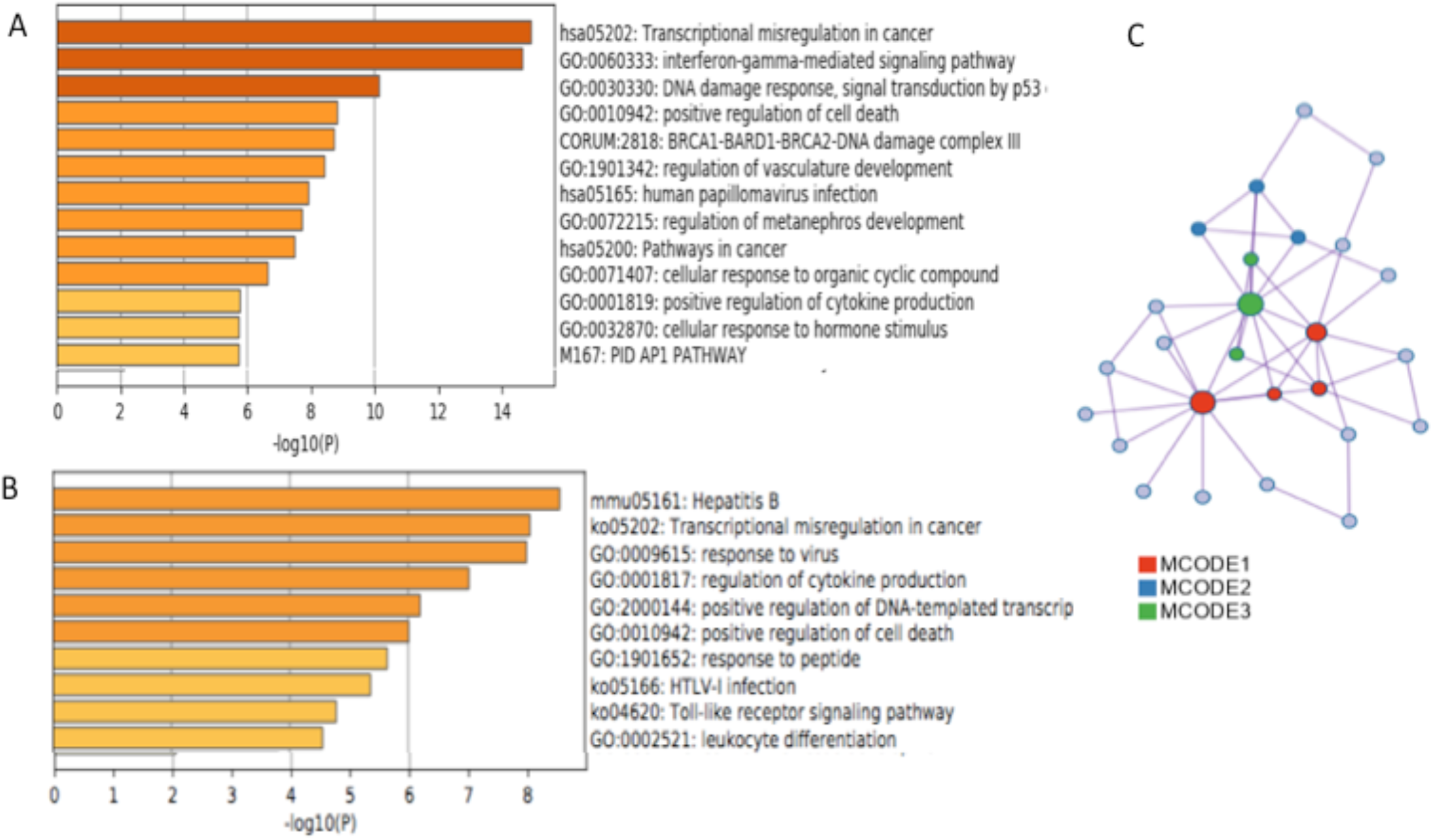
Annotation of biological functions and signal transduction pathways in human PBMC and mouse immune cells using Metascape software tools (A) Biological functions and signaling pathway gene ontology (GO) terms associated with transcription factors of human PBMC were ranked by significance (B) Biological functions and signaling pathway gene ontology (GO) terms associated with transcription factors of mouse immune cells were ranked by significance (C) Molecular complex detection (MCODE) algorithm detection of modules associated with transcription factors induced by IFN-β in human PBMC.

**Figure 2.**
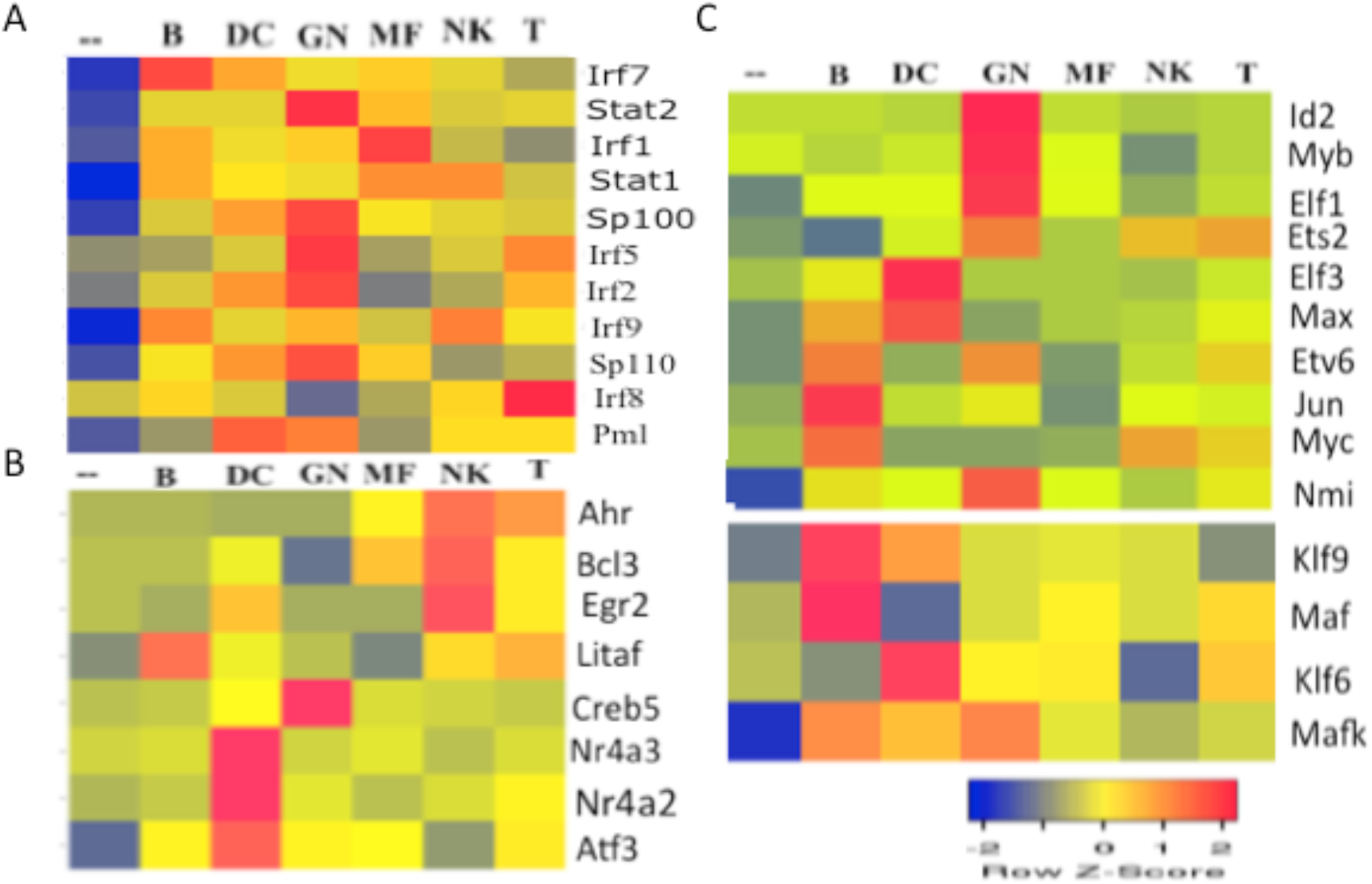
Functional organization of transcription factor gene expression in the transcriptome induced by IFN-α in mouse immune cell types (A-C) Transcription factor sub-networks of antiviral, immune modulation, and cell growth were retrieved from microarray datasets of mouse immune cells treated with IFN-α for two hours. Expression in B-lymphocytes (B), dendritic cells (DC), granulocytes (GN), macrophage (MF), natural killer cells (NK), and T-lymphocytes (T) were shown. Cluster analysis was performed using Heatmapper software tools.

**Figure 3.**
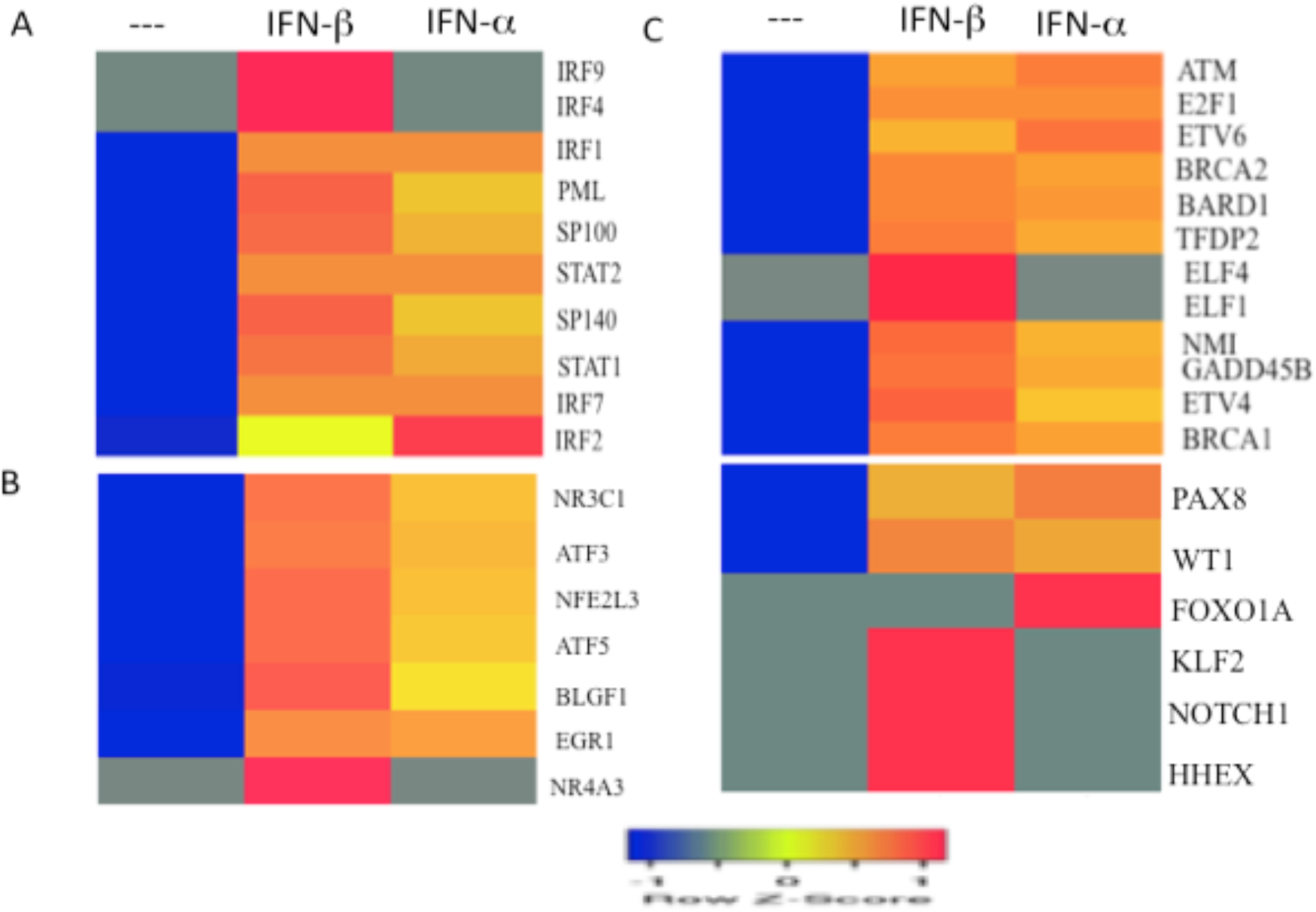
Functional organization of transcription factors induced by IFN-α or IFN-β in the human PBMC (A-C) Transcription factor sub-networks of antiviral, immune modulation, and cell growth was retrieved from microarray dataset of human PBMC treated with IFN-α or IFN-β for 4 hours. Cluster analysis was performed using Heatmapper software tools.

### 3.2 Time-course analysis of Type I Interferon signaling

Most of the gene expression profiling studies in response to type I IFN have some technical limitations (5). These studies were performed in transformed cells or cells cultured in the presence of high serum or in a limited number of cell types such as fibroblasts or epithelial cells. In addition, cells were subjected to prolonged interferon treatment of several hours to capture the maximum range of gene expression levels. Furthermore, antiviral response genes were overrepresented while transcription factor and growth-regulated genes were under-represented on many custom-designed arrays of interferon-regulated gene expression. These studies may have missed earlier dynamic changes in transcriptional factor mRNA levels and cell growth-related gene expression. Interrogation of gene expression datasets of early time points after type I Interferon treatment revealed that several transcription factor mRNA were expressed after 1-4 hour treatment of human PBMC and mouse B cells. (Figure 4A and 4B). Transcription factors involved in antiviral response including Stat1, Stat2, Irf1, Irf2, and Irf7 mRNA levels were rapidly induced in both PBMC and B cells (Figure 4). An intact type I IFN response is necessary to inhibit viral replication in cells, control the tissue restriction of virus, and increase the production of type I IFN acting as a feedback loop (4,36). A large number of type I interferon regulated genes like 2,5-oligoadenylate synthetase (OAS), Mx, PKR, and Rnase L implicated in antiviral defense were induced rapidly after infection. Mutations of Stat1 have been characterized in human populations. Patients with heterozygous Stat1 mutations show susceptibility to mycobacteria but not the viral disease. Interferon-induced Stat1 homodimer or gamma-activated factor (GAF) formation was diminished, while the response to the ISGF-3 complex was normal in cells derived from these patients (37). The clinical and cellular phenotypes of these patients suggest that the response to mycobacteria was mediated by Stat1 homo-dimer and the antiviral immunity is dependent on ISGF-3 formation. Interestingly, transcription factors and growth-related genes such as Myc, Jun, and Schlafen (Sfn) family members were rapidly induced by type I Interferon in B cells (Figure 4B). Schlafens regulate immune cell proliferation, differentiation, and restricting virus replication (38). However, Schlafens has no DNA binding activity or direct role in transcriptional regulation.

**Figure 4.**
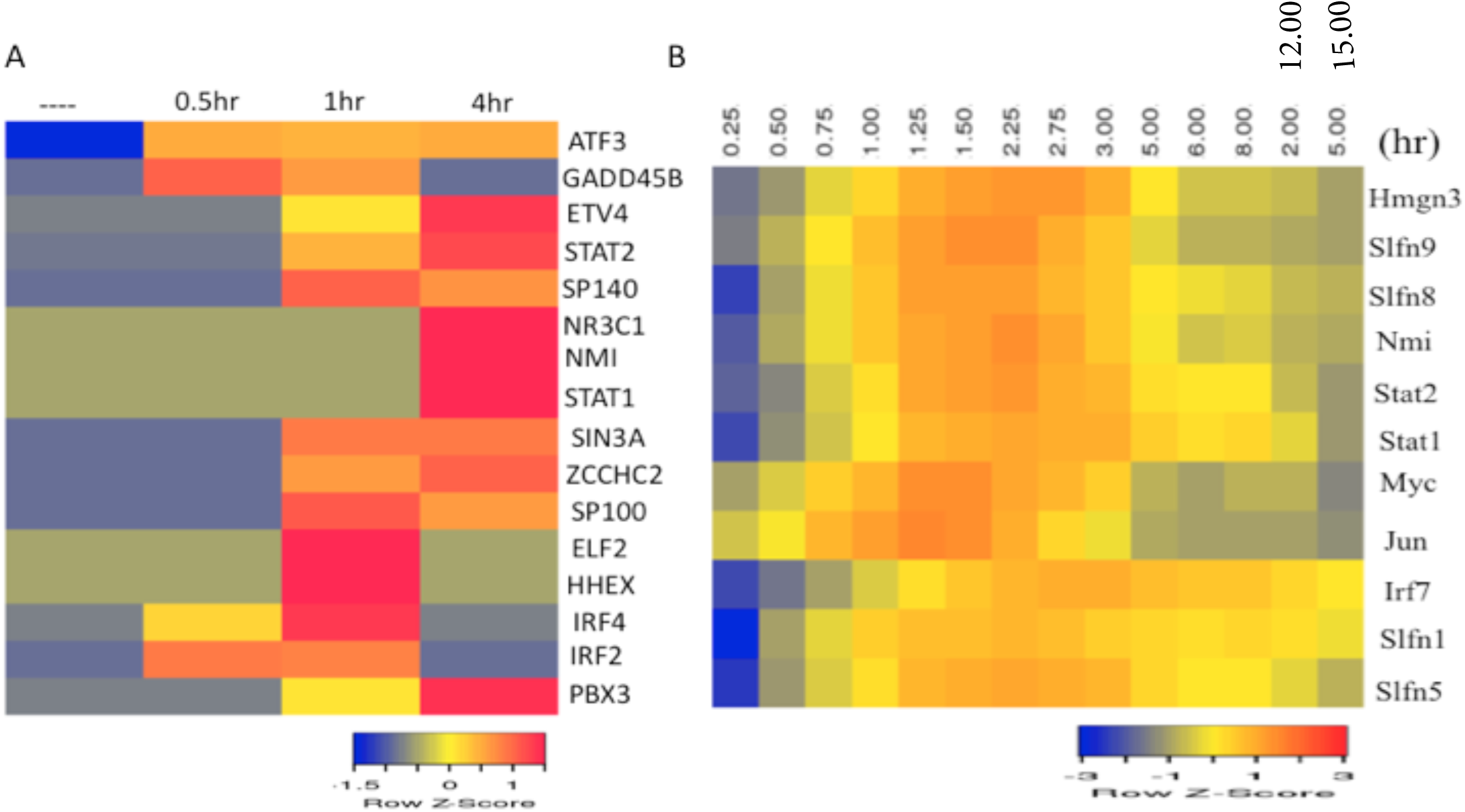
Time-course analysis of type I interferon-induced transcription factor (TF) mRNA levels in human and mouse immune cells (A) Human PBMC treated with IFN-β for 0.5-4 hours (B) Mouse B lymphocyte cells treated with IFN-α for 0.25 to 6 hours. The gene expression data were retrieved from different sources as described in materials and methods section. Cluster analysis of gene expression across the arrays was accomplished using the Heatmapper software.

### 3.3 Protein-protein interaction network and functional connectivity of signaling hubs in type I Interferon signaling

Transcriptional factors bind to regulatory elements such as promoters and enhancers located upstream of the gene transcription start site and act as platforms for the assembly of co-regulators and with general transcriptional factors regulate eukaryotic gene expression (39). Co-operative protein-protein interactions play an important role in the functional diversity of signal transduction pathways and biological functions of multiple cell types (40). Stat1 plays a major role in the transcriptional regulation by type I and type II interferons (1-3). Stat1 is regulated by post-translational modifications such as tyrosine and serine phosphorylation and interacts with a large number of general and specific transcription factors. Stat1 requires distinct components of the co-activator complex, including the CREB-binding protein family of transcriptional activators (CBP/p300), and p300/CBP-associated factor (p/CAF) for distinct platform assembly and histone acetyltransferase (HAT) activity to activate interferon-stimulated gene transcription (41). Two contact regions between Stat1 N-terminal and C-terminal regions and CBP/p300 were identified in interferon signaling (42). RNA polymerase II and associated general transcription factors such as upstream stimulatory factors (USF) and TATA box binding protein (TBP) play an important role in transcriptional initiation (43). In contrast to Stat1, the role of inducible transcription factors in type I interferon signaling is not well understood. It is possible to construct potential protein interaction networks of inducible transcription factors of distinct functional categories in type I Interferon signaling using the data from protein-interaction databases (33-35,44). As an example, antiviral transcription factor interactions in the STRING database was shown as a graph with proteins or nodes as ovals and protein interactions as edges (Figure 5A). Network analysis in TRUUST database revealed extensive interactions between Stats, IRF, and Sp family of transcription factors (45, data not shown). A major advantage of this approach is that it allows tools developed for social network analysis such as centrality measures to be applied to understand the organization of protein interaction networks and critical nodes. These graph theory algorithms reveal the importance of any particular node to the entire network. Some of the well-known centrality methods include degree centrality that measures the node connectivity or the total number of inbound and outbound links. In contrast, between centrality identifies the nodes that are bridges on the shortest path between other nodes. Furthermore, closeness centrality measures the closeness of any node to other nodes in the network (46). Applying all these measures facilitated the ranking of each node in the antiviral sub-network (Table 2). These calculations have shown that Stat1, Stat2, and Irf1 were the highest-ranked nodes in the antiviral sub-network. In contrast, Irf5 and Sp110 were the lowest-ranked nodes in the sub-network. Furthermore, transcription factors involved in a related biologic function often share common protein interaction partners and target genes as revealed by pairwise analysis of transcription factor hubs involved in innate and adaptive immunity (25). Implementing the pairwise analysis of common protein interaction partners of Stat1 and other hubs in antiviral sub-network in Biogrid database revealed that Rela and Stat3 as additional members of the extended network (data not shown). Although Stat1 was included as a hub in the antiviral sub-network, it is possible to include Stat1 in immune modulation or cell growth sub-network (Figure 5B). Such a hub can be designated as an authority among hubs. In social network analysis hubs and authority designations were developed in the context of hyperlink-induced topic search (47). Type I Interferon is often released simultaneously with pro-inflammatory cytokines such as TNF-α, IL-1. IL-6 by immune cells in response to respiratory virus infection resulting in the cross-talk between Stat1 and other transcription factors such Stat3, Rela, and Jun and fine-tuning of pathogen response (48). The protein interaction module of Stat1, Stat3, Rela, and Jun was shown to be a characteristic feature of innate and adaptive immunity (25)

**Table 2.**
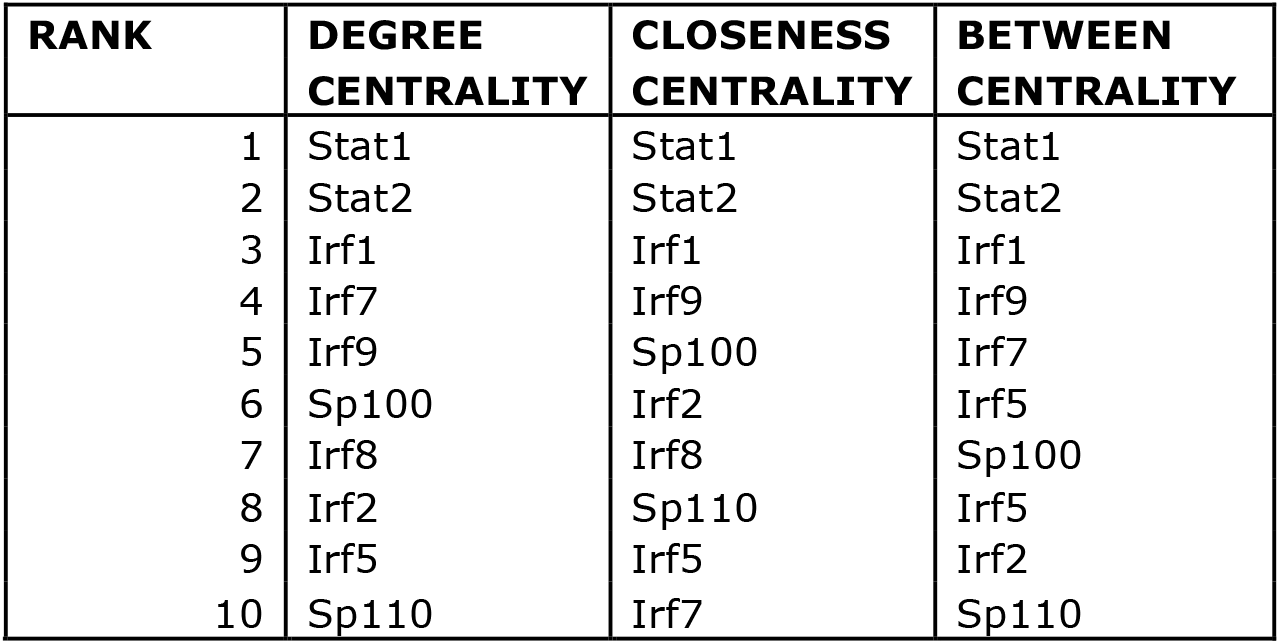
Centrality measures for ranking nodes in antiviral sub-network

| **RANK** | **DEGREE CENTRALITY** | **CLOSENESS CENTRALITY** | **BETWEEN CENTRALITY** |
| --- | --- | --- | --- |
| 1 | Stat1 | Stat1 | Stat1 |
| 2 | Stat2 | Stat2 | Stat2 |
| 3 | Irf1 | Irf1 | Irf1 |
| 4 | Irf7 | Irf9 | Irf9 |
| 5 | Irf9 | Sp100 | Irf7 |
| 6 | Sp100 | Irf2 | Irf5 |
| 7 | Irf8 | Irf8 | Sp100 |
| 8 | Irf2 | Sp110 | Irf5 |
| 9 | Irf5 | Irf5 | Irf2 |
| 10 | Sp110 | Irf7 | Sp110 |

**Figure 5.**
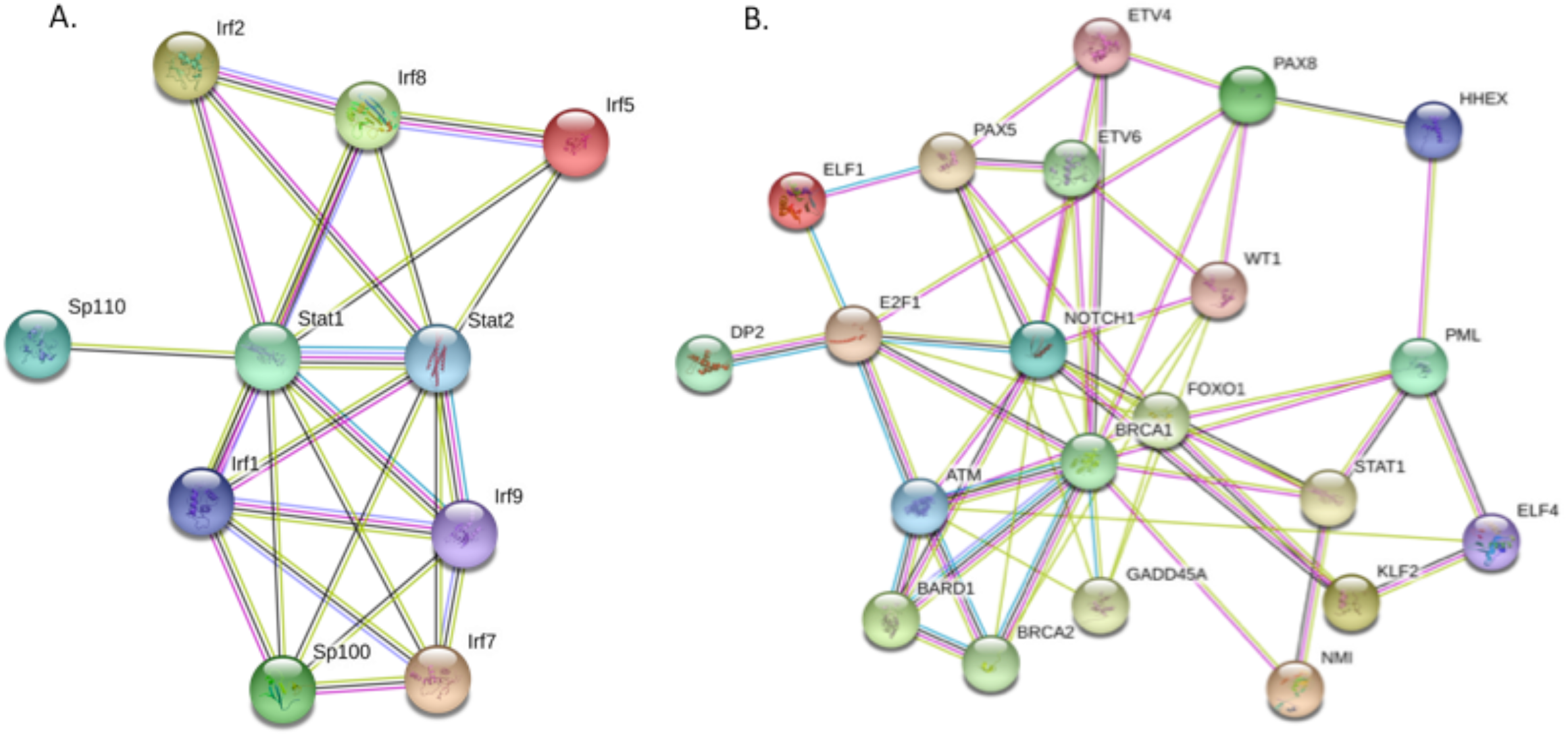
Protein-protein interactions of transcription factor sub-networks induced by type I interferon in mouse and human immune cells represented in the STRING database (A) Antiviral sub-network of mouse immune cells induced by type I interferon (B) Cell growth sub-network of human PBMC induced by type I interferon. Transcription factors were represented by ovals and protein interactions by edges, respectively.

### 3.4 Regulation of Core Type I Interferon signaling in immune cells

Phylogenetic analysis of type I Interferon regulated gene expression across multiple species provided novel insights into the origin and evolution of innate immunity (49). Many components of innate immunity and interferon signaling like pathogen sensors, transducers, transcription factors such as interferon regulatory factors (IRF), and target genes were already present and functional before the origin of the type I interferon signaling system in fishes (50). Microarray analysis of peripheral blood monocytic cells (PBMC) treated with type I interferon across multiple vertebrate species revealed a set of 62 highly conserved interferon-stimulated genes (ISG) designated as core ISG involved in innate immunity (51). The functional sub-categories of these genes include antiviral defense, antigen presentation, the immunoproteasome, ubiquitin-modification, cell signaling, and apoptosis. This list includes RNA-sensing pathogen recognition receptors such as Ifih (Mda5), Dhx58 (Lgp2), Rig-I (Ddx58), and the adaptor molecule Myd88. The highest level of Core ISG expression was observed in granulocytes compared with other immune cells and includes genes involved in antiviral (Oas, Mx, Eif2ak2), cell growth and apoptosis (Tnfsf10, Parp9, Parp14, Casp8), and MHC class I antigen presentation (Figure 6 and data not shown). The group of inducible transcription factor mRNA in core type I Interferon signaling was dominated by mediators of antiviral response such as Stat1, Stat2, Irf1, Irf7, Irf9, Pml, and Sp110. Many of the core antiviral response transcription factors were enriched in granulocytes and interact with each other to form a tight protein interaction network (Figure 7A and 7B). Consistent with the results, promoter scanning of core ISG by P-Scan program revealed a highly significant representation of binding sites for Stat1, Stat2 and IRF family members (data not shown). These studies revealed that the evolutionarily conserved core ISG primary function was the antiviral response and other biological functions such as immune modulation and cell growth were elaborated later in the evolution. Novel inducible core transcription factors detected in this functional screen include two zinc finger proteins Zcc3hav1, Zcchc2 involved in the regulation of antiviral response and cell growth, respectively (52,53).

**Figure 6.**
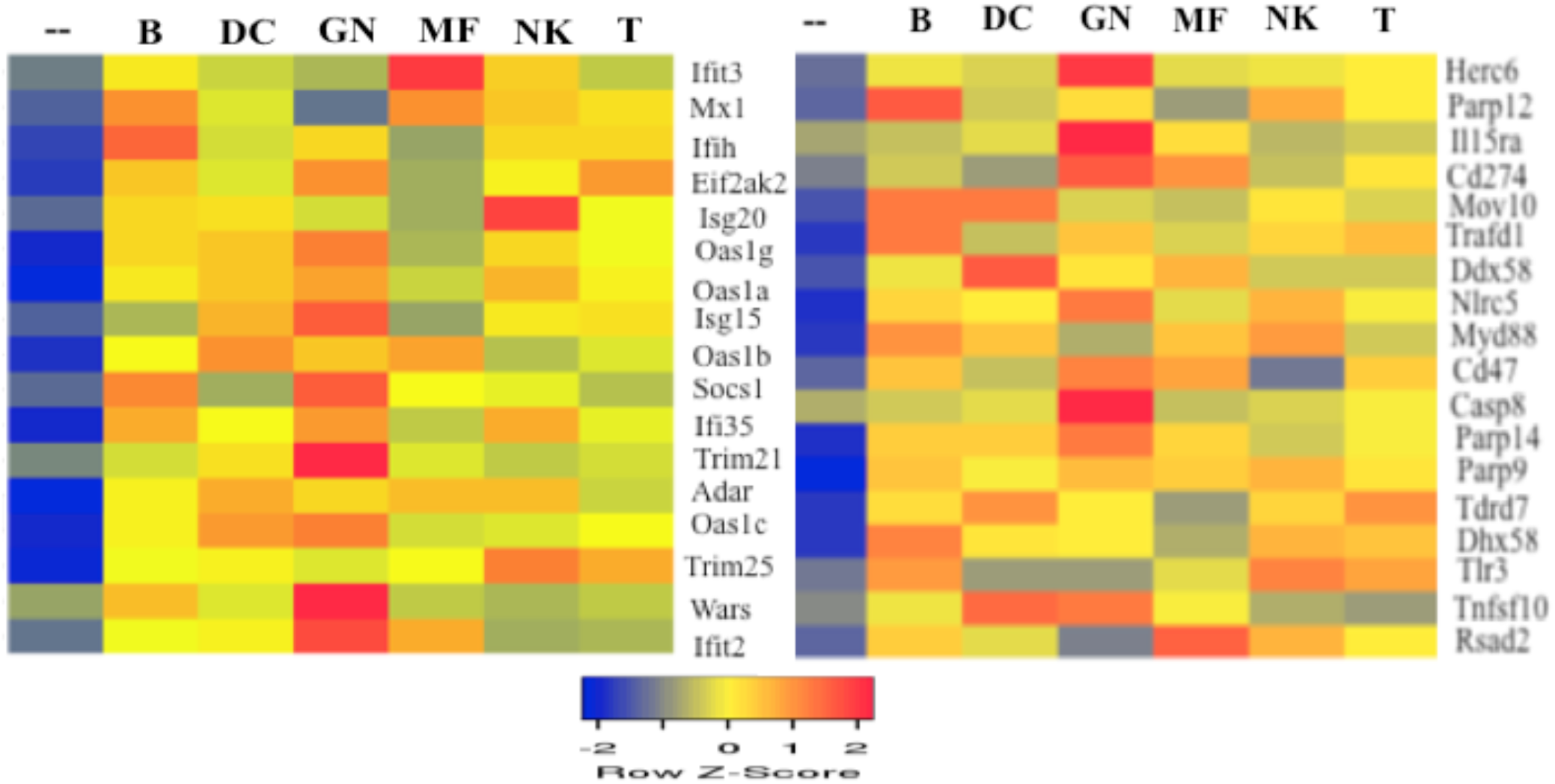
Core interferon-stimulated gene expression (core ISG) in mouse immune cells. Cluster analysis of core ISG expression levels in B-lymphocytes (B), dendritic cells (DC), granulocytes (GN), macrophage cells (MF), natural killer cells (NK), and T-lymphocytes (T) were shown.

**Figure 7.**
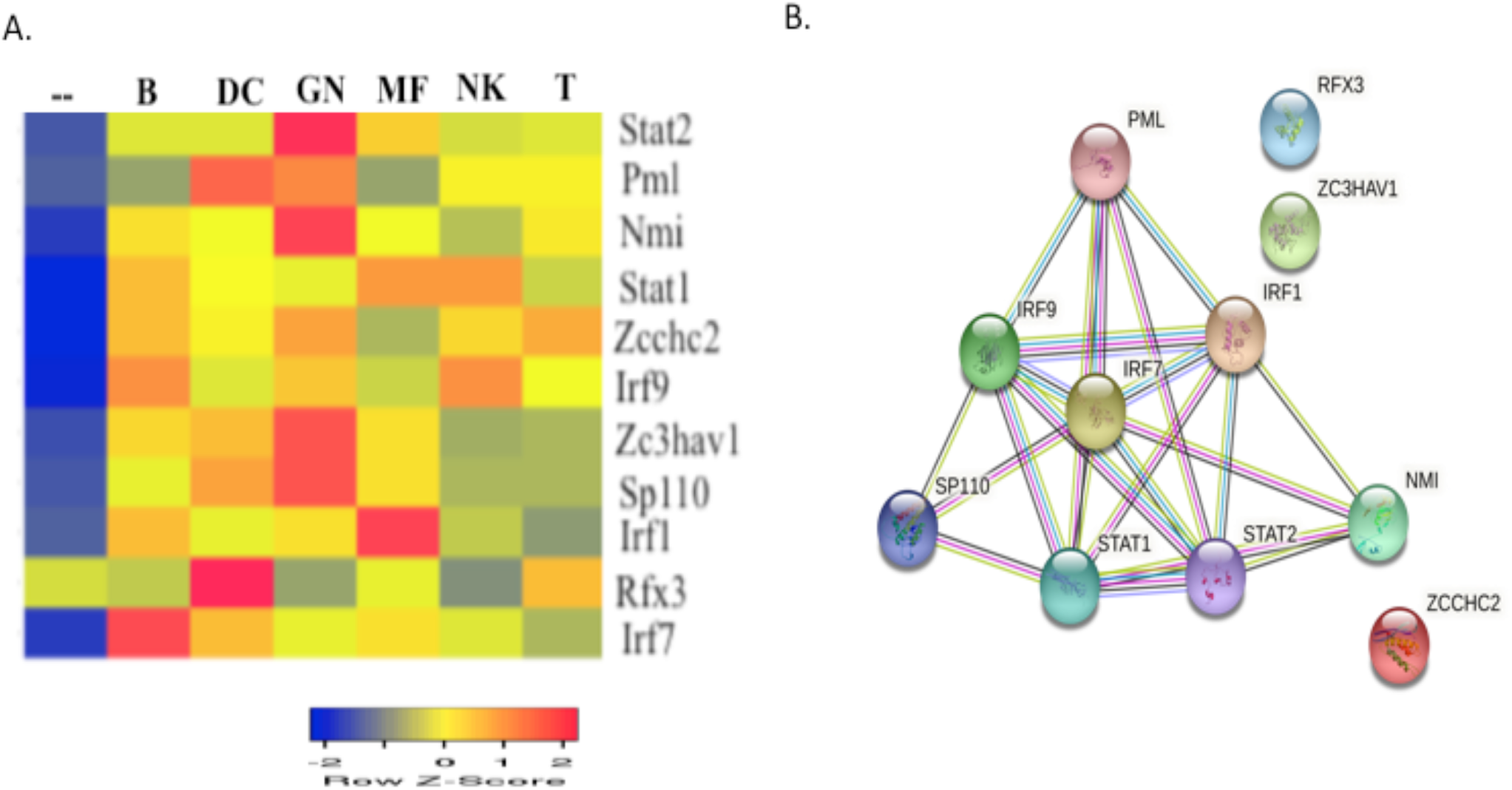
Expression levels of core interferon-inducible transcription factor (TF) mRNA and protein-protein interaction network in mouse immune cells (A) Cluster analysis of interferon-inducible transcription factor expression levels in B-lymphocytes (B), dendritic cells (DC), granulocytes (GN), macrophage cells (MF), natural killer cells (NK), and T-lymphocytes (T) were shown. (B) Protein interaction network of transcription factors in STRING database was shown.

### 3.5 Role of tyrosine kinase Tyk2 and alternative signaling pathways for inducible transcription factors in Type I Interferon signaling

Type I interferon signaling involves the binding of IFN α/β to its receptor (IFNAR) and activation of receptor-associated protein tyrosine kinases Jak1 and Tyk2 by auto and trans-phosphorylation (1,3,54). These kinases are involved in the phosphorylation of the receptor on specific tyrosine residues (Y337, Y512) that serve as docking sites for the recruitment and subsequent activation of Stat1 and Stat2 (55). Tyk2-null cells are impaired in the activation of Stat1 and Stat2, and ISRE-mediated gene expression (56,57). Furthermore, the induction of several transcription factor mRNA by IFN-α was attenuated in Tyk2-null B-cells suggesting that activation of Tyk2 is required for Stat-dependent and –independent pathways of signaling (Figure 8A). In addition to Jak kinases, IFNAR also activates alternative pathways involving multiple kinases in parallel such as extracellular signal-regulated kinases (Erk1/2), p38 mitogen-activated protein kinase (p38 MAPK), phosphatidylinositol-3 kinase (PI-3K), Akt serine kinase, and protein kinase C-delta (3,67,68). Jak1 and Tyk2 interact with a large number of proteins as documented in protein-interaction databases and common protein interaction partners of Jak1 and Tyk2 are potential candidates for alternative pathways of type I interferon signaling (25).

**Figure 8.**
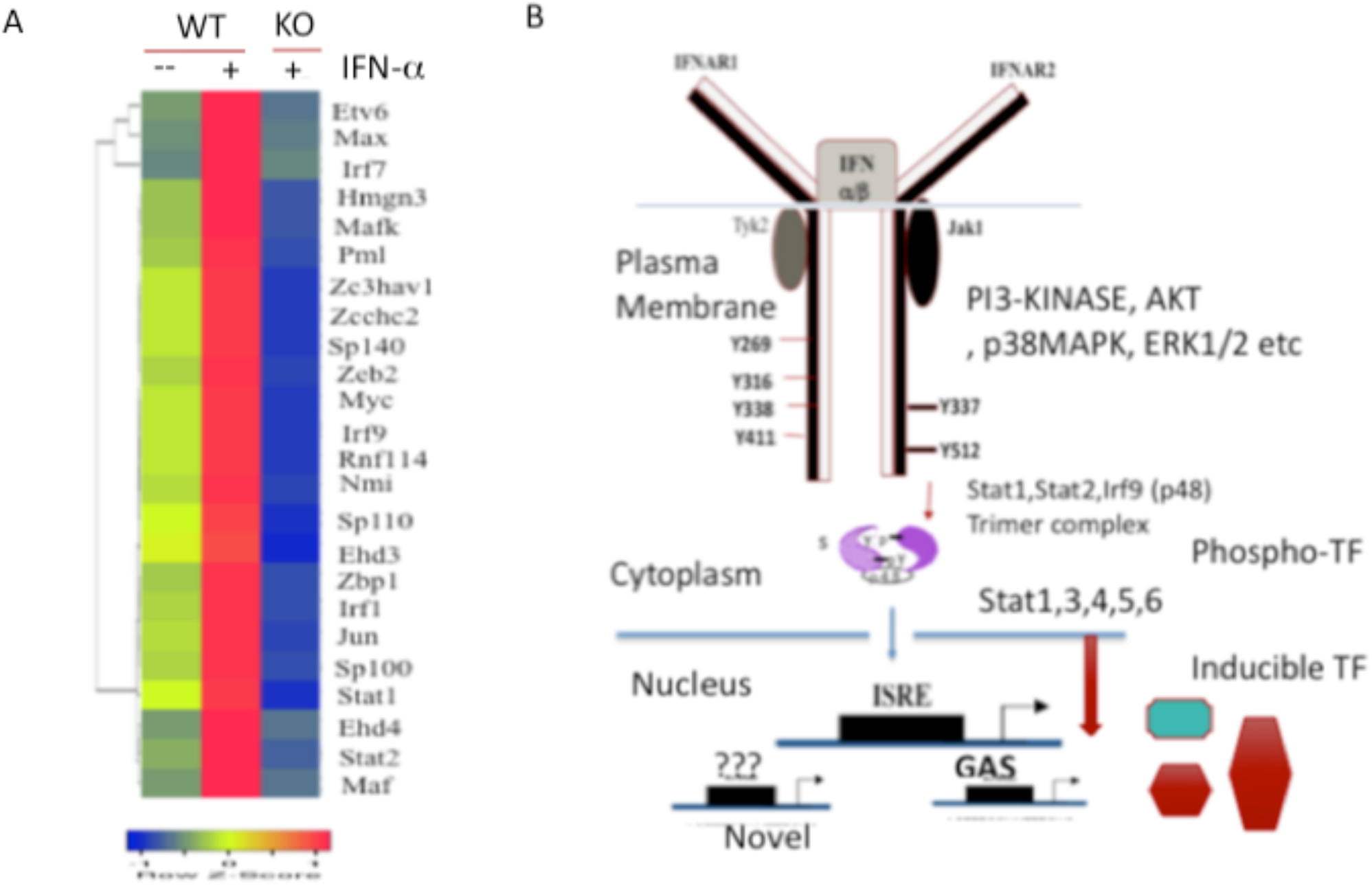
Role of tyrosine kinase Tyk2 and signaling pathways in phosphorylation–dependent and inducible transcription factors in Type I Interferon signaling (A) Expression levels of transcription factor mRNA in wild-type B cells untreated (control), wild-type B cells (Wt), and Tyk2 knock-out B cells (KO) treated with IFN-α for 2 hours (B) A model for type I interferon signaling pathways involving rapid phosphorylation-dependent activation of Stat transcription factors (0.5 hr) and gradual increase of inducible transcription factors (1-4 hr) in transcriptional regulation

IFN-α/β has been shown to activate multiple Stat members such as Stat3, Stat4, Stat5, and Stat6 in different cell types as demonstrated by tyrosine phosphorylation and DNA binding assays. Activation of these Stat members in the regulating of antiviral response, cell proliferation, and differentiation has been observed in different cell types (58-63). IFN-α/β activated Stat3 in RAW264.7 macrophages and in primary B-lymphocytes (data not shown). Catalytically active Tyk2 was required for IFN-β mediated tyrosine phosphorylation and activation of Stat3 (64). Stat3 regulates a large number of target genes including Myc, Jun and Schafen2 that are rapidly induced by IFN-α in B cells (Figure 2). Positive and negative regulation of type I interferon signaling by Stat3 was reported. Two patients with a homozygous mutation of the Stat1 gene were identified (65). Both individuals suffered from mycobacterial disease but died of viral disease. IFN-α treatment of B-lymphocyte cell lines derived from these patients resulted in the tyrosine phosphorylation of Stat3, but no activation of ISGF-3 was observed. Furthermore, IFN-α induced Stat3 target genes such as Suppressor of cytokine signaling -3 (SOCS-3) in these Stat1-deficient cell lines indicating that IFN-α/β stimulated Stat1-independent signaling in human cells. Stat3 activation may have a negative role in type I interferon signaling by antagonizing multiple steps in ISGF3 transcriptional activity (66). The physiological significance of activation of additional members of the Stat family in the induction of transcription factor mRNA by IFN α/β remains to be elucidated. A schematic model of type I interferon signaling involving canonical Jak1 and Tyk2 leading to the rapid activation of ISGF3 and additional Stat family members by tyrosine phosphorylation was shown (Figure 8B). The members of the Stat family of transcription factors share structure, activation mechanism and have distinct binding affinities for cis-elements in the gene promoter leading to the regulation of a different set of genes. Different combinations of Stats may be activated in distinct cell types resulting in a variety of biological effects. The cytoplasmic domain of IFNAR contains several tyrosine phosphorylation sites that are not involved in Stat activation (69). The role of these additional IFNAR phosphorylation sites and activation of multiple kinases in type I interferon signaling and inducible transcription factor mRNA expression remains to be elucidated (Figure 9B).

**Figure 9.**
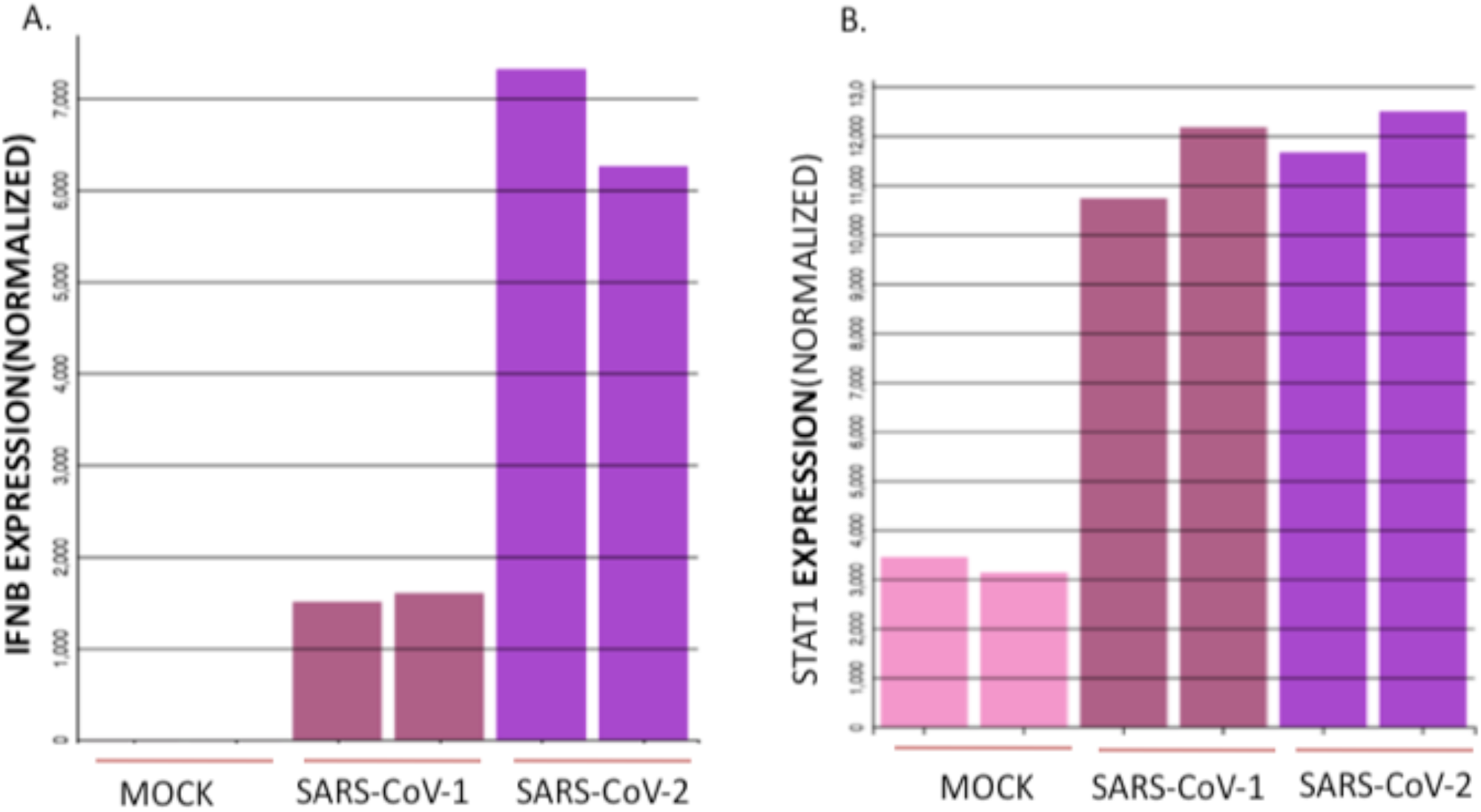
Regulation of IFNB1 and STAT1 mRNA levels by corona viruses in Calu-3 lung epithelial cells (A) Calu-3 cells were mock treated or infected with SARS-CoV-1 or SARS-COV-2 for 24 hours. IFNB1 mRNA expression values normalized by DEseq2 were shown (B) STAT1 mRNA expression values normalized by DEseq2 under the same experimental conditions were shown. Data from two independent samples for each condition were shown.

### 3.6 Profiling inducible transcription factors of Type I Interferon signaling in lung epithelial cells after SARS-CoV-2 infection and in the tissues of COVID-19 patients

Respiratory virus infection of lung epithelial cells results in type I interferon production, which then acts in an autocrine or paracrine manner to stimulate interferon-stimulated gene expression (16,17). Genomic analysis of critically ill COVID-19 patients revealed deficiency or alterations in type I interferon signaling (21-23). Imbalanced cytokine and interferon responses in COVID-19 patients were described (27,70,71). Autoantibodies to type I interferon and bacterial products were also detected in the tissue samples of COVID-19 patients (22,70). Furthermore, the differential expression of ISG was correlated with lung damage and survival in COVID -19 patients (72). Respiratory virus infection leads to the immunopathology of the lung mediated by the direct effect of the virus as well as imbalanced host immune response (19,73,74). Gene expression data of human lung epithelial cells infected with SARS-CoV-2 as well as healthy and COVID 19 tissue samples were published recently, making it possible to study type I interferon regulation of inducible transcription factor gene expression (27,28). Interrogation of microarray datasets revealed that interferon beta (IFN-β) and STAT1 mRNA levels were significantly increased 24 hours after infection with SARS-CoV-1 or SARS-CoV-2 in human Calu-3 lung epithelial cells (Figure 9). Consistent with these results, type I interferon-inducible transcription factor mRNA levels involved in biological functions such as antiviral response, inflammation, and cell growth were significantly enhanced in Calu-3 cells infected with SARS-CoV-1 and SARS-CoV-2 (Figure 10). In contrast, SARS-CoV-2 infection of human A549 lung type II cells did not induce IFN-β mRNA (Figure 11A). It has been shown that SARS-CoV-2 entry into lung epithelial cells required the expression of virus entry receptor Angiotensin-converting enzyme 2 or ACE2 (75). Infection of A549 cells expressing ACE2 with SARS-CoV-2 rescued in the inducible expression of IFN-β and STAT1 mRNA within 24 hours (Figure 11A and Figure 11B). Furthermore, expression of type I interferon-inducible transcription factor mRNA levels involved in biological functions such as antiviral response, immune modulation, and cell growth were significantly enhanced in A549 cells expressing ACE2 receptor (Figure 12). A high level of interferon-stimulated gene expression (ISG-high) in patients was correlated with high viral titers and high levels of cytokines and limited lung damage. In contrast, a low level of interferon-stimulated gene expression (ISG-low) in COVID-19 patients was correlated with lower viral loads, extensive lung damage with the presence of activated CD8^+^ T cells, and macrophages (72). ISG-high expressing patients died significantly earlier after hospitalization than ISG-low expressing patients suggesting that type I interferon signaling intensity may be used as a prognostic feature of survival in COVID-19 patients (72). Stat1 mRNA levels were increased in COVID 19 lung compared with healthy lung tissue samples (Figure 13A). Furthermore, inducible transcription factors involved in antiviral response were up-regulated and the transcription factors involved in inflammation and cell growth were down-regulated in COVID 19 lung compared with healthy lung samples (Figure 13B). The peripheral blood monocytic cells (PBMC) composed of several distinct cell types and the distribution of changes in gene expression varied significantly in different cell types, in response to type I interferon (Figures 2 and 3). Expression of the inducible transcription factors of the antiviral, cell growth, and inflammation sub-networks were specifically enhanced in neutrophils of COVID-19 patients, compared with healthy subjects (Figure 14A-C). High-level expression of a subset of type I interferon-stimulated genes in tissue samples of COVID-19 patients was observed including a gene signature of STAT1, STAT2, TNFSF10, S100A8, S100A9, and S100A12. This gene signature was similar to the up-regulated gene expression profile observed in patients with the autoimmune disease known as Sjogren’s syndrome, characterized by systemic inflammation (76,77). This gene expression signature was highly expressed in both the lungs and PBMC of COVID19 patients (Figure 13C and Figure 14B). Elevated expression of inflammatory markers such as EGR1, TNFSF10, TNFSF14, S100A8, and S100A9 was reported in COVID-19 patients (26,70,78). Members of the TNF superfamily such as TNFSF10 (TRAIL) and TNFSF14 (LIGHT) were implicated in apoptosis and inflammation (79). The S100 family members include S100A8. S100A9 and S100A12 are intracellular calcium-binding proteins that are released into extracellular space and function as damage-associated molecular pattern molecules (DAMPS) involved in tissue repair (80). These studies demonstrate that IFN-a/β mediated induction of three distinct transcription factor sub-networks in human and mouse immune cells. Furthermore, differential regulation of these transcription factor sub-networks was observed in human lung epithelial cell lines in response to SARS-CoV-2 infection, and in the tissue samples of healthy and COVID-19 patients.

**Figure 10.**
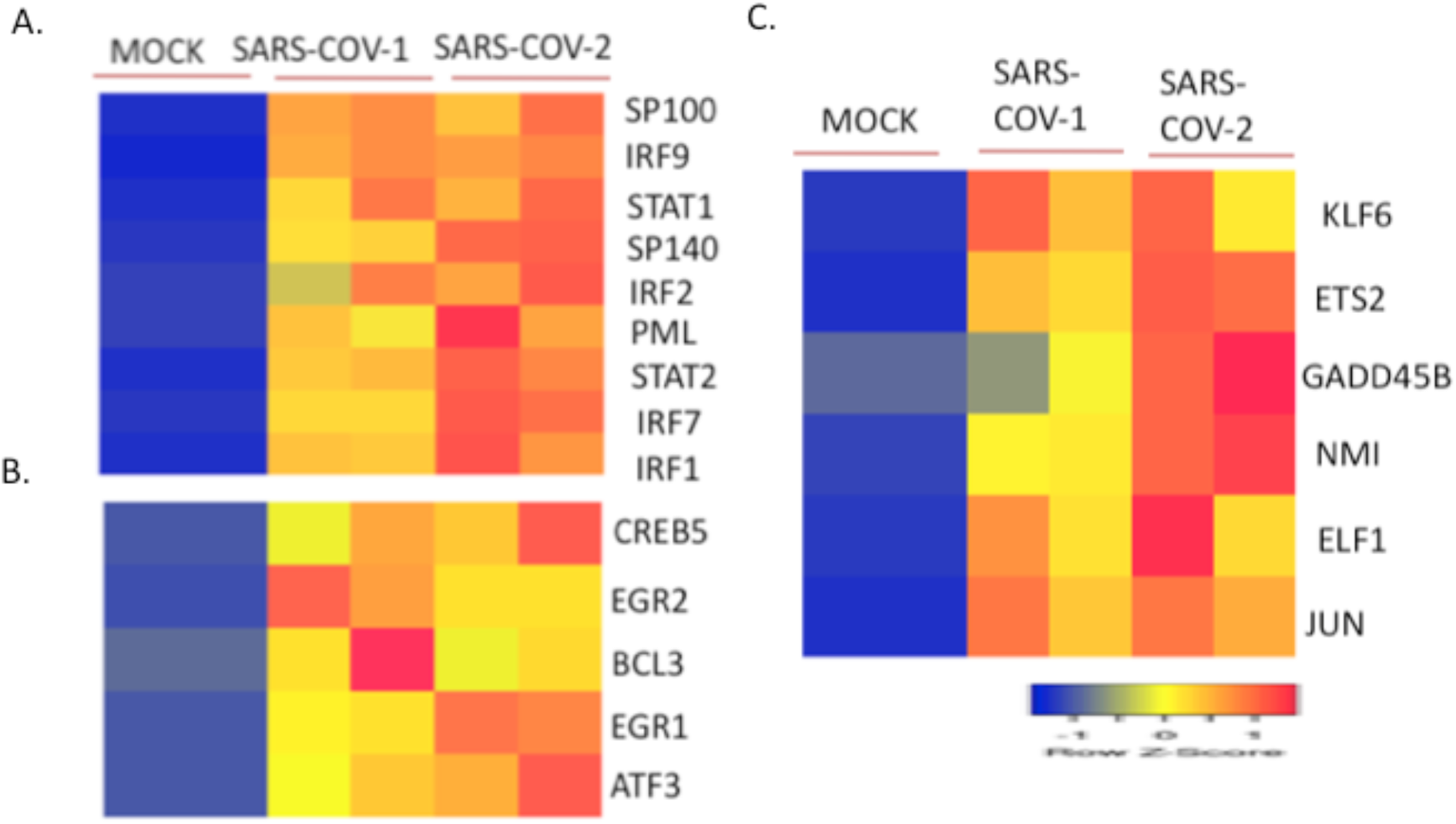
Regulation of transcription factor mRNA levels of type I interferon sub-networks by coronaviruses in Calu-3 lung epithelial cells. Calu-3 cells were mock treated or infected with SARS-CoV-1 or SARS-CoV-2 for 24 hours (A) Transcription factors of the antiviral sub-network were shown (B) Transcription factors of the immune modulation sub-network were shown (C) Transcription factors of the cell growth sub-network were shown. Gene expression data from two independent samples for each condition were shown.

**Figure 11.**
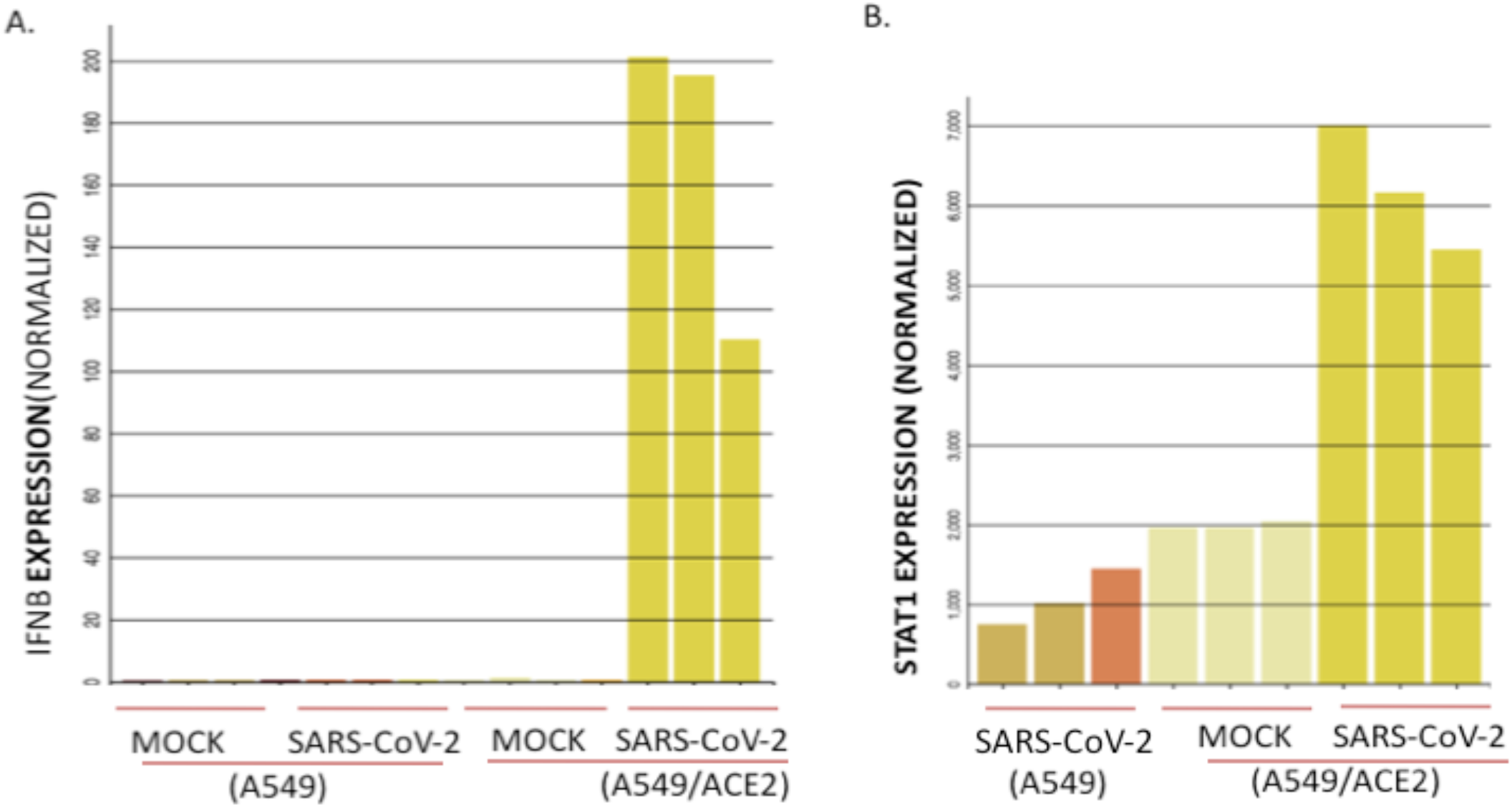
Regulation of IFNB1 and STAT1 mRNA levels by SARS-CoV-2 in A549 lung type II cells (A) A549 or A549 cells expressing ACE2 receptor were mock treated or infected with SARS-CoV-2 virus for 24 hours. IFNB1 mRNA expression values were normalized by DEseq2 (B) STAT1 mRNA expression values were normalized by DEseq2 under the same conditions were shown. Data from three independent samples for each condition were shown.

**Figure 12.**
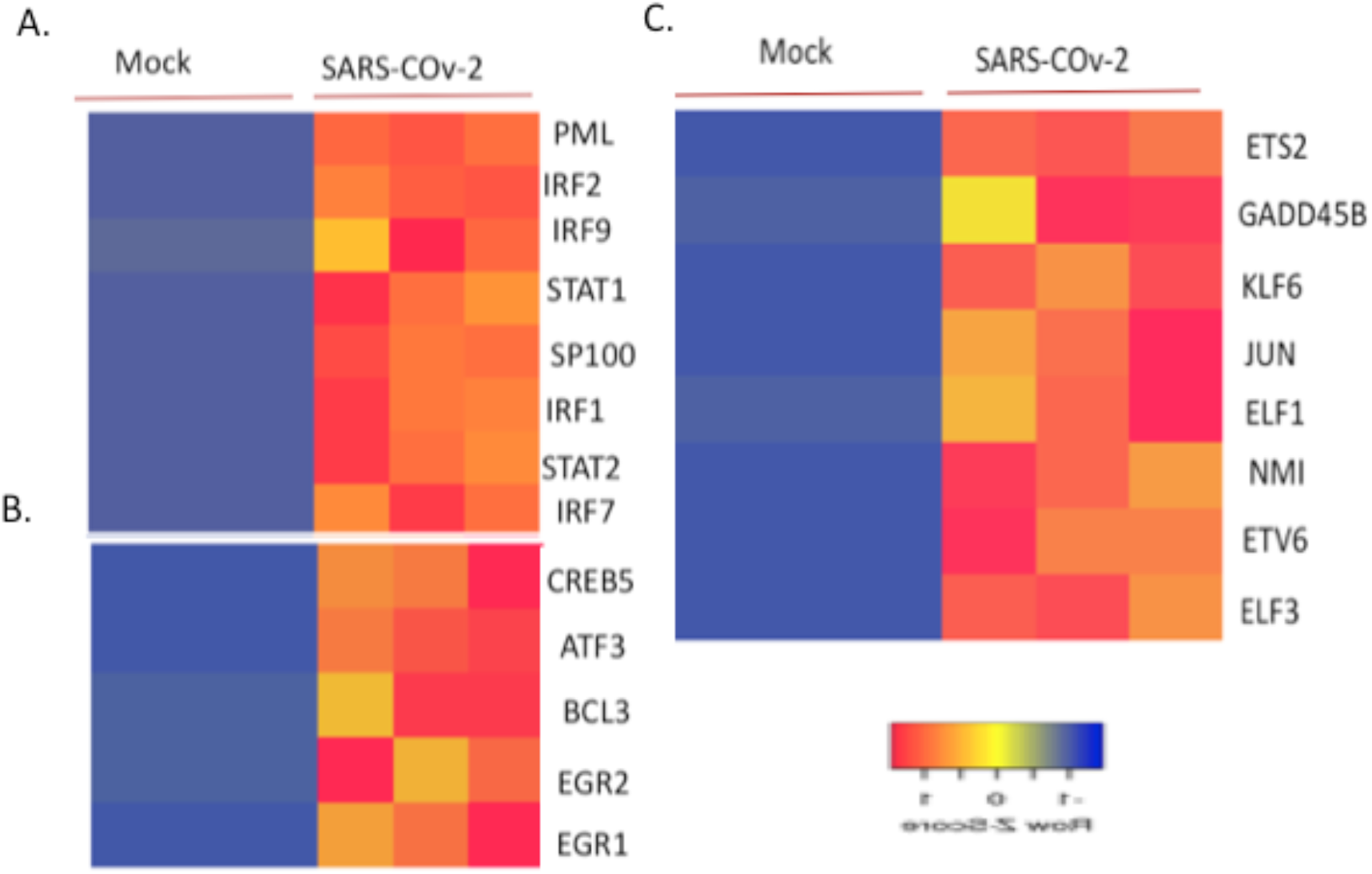
Regulation of transcription factor mRNA of type I interferon sub-networks by SARS CoV-2 in A549 cells expressing ACE2 receptor (A) A549 cells expressing virus entry receptor ACE2 were mock infected or infected with the virus SARS-CoV-2 for 24 hours. Transcription factors of the antiviral sub-network were shown (B) Transcription factors of the immune modulation network under the same conditions were shown (C) Transcription factors of the cell growth sub-network under the same conditions were shown. Three independent samples for each condition were shown.

**Figure 13.**
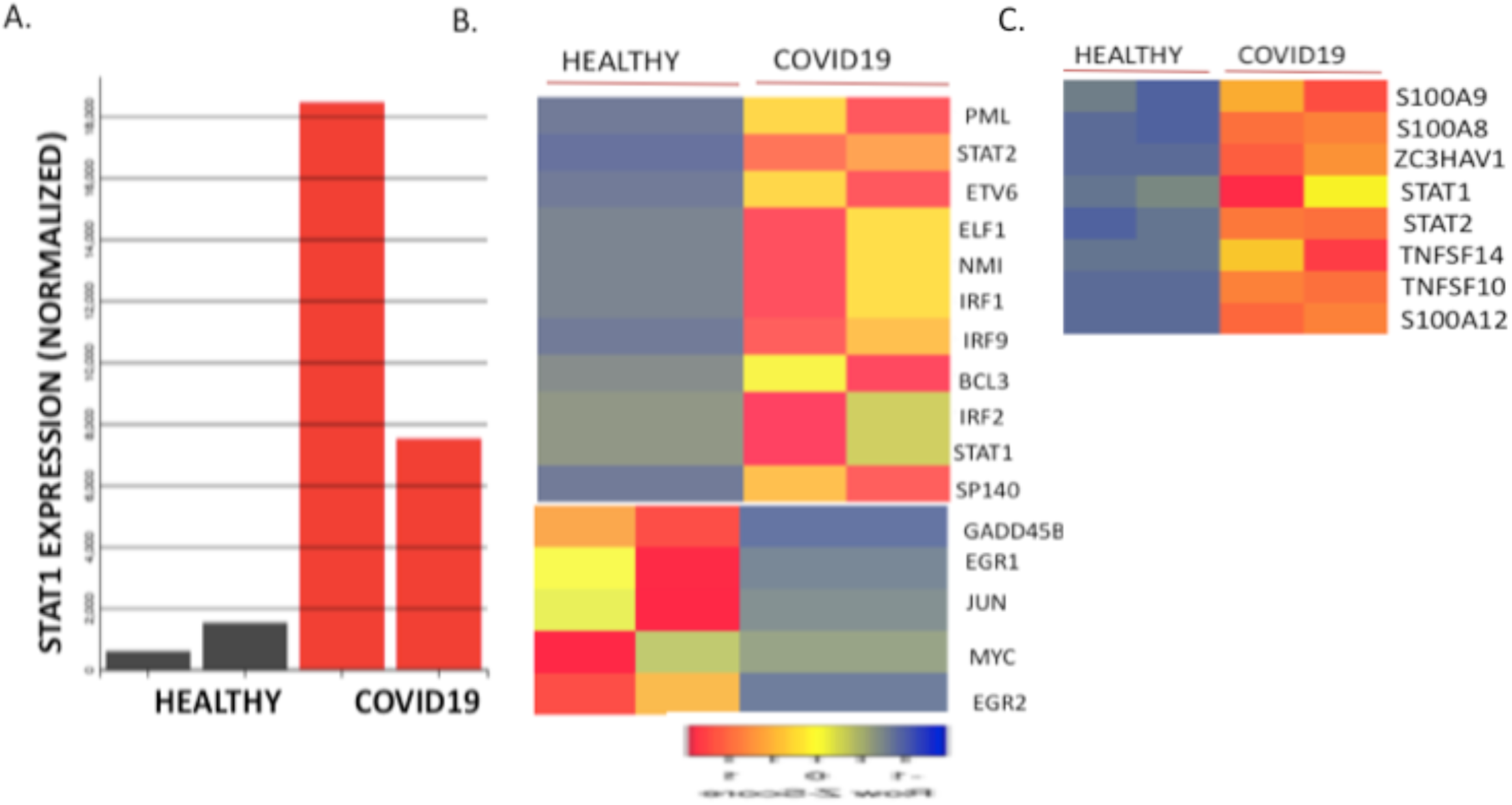
Regulation of transcription factor and inflammatory biomarker mRNA levels by type I interferon signaling in the lung biopsies of healthy and COVID19 patients (A) Relative mRNA expression levels of STAT1 in the lung biopsies of healthy and COVID19 patients (B) Transcription factors of the antiviral, immune modulation and cell growth sub-networks in the lung biopsies of healthy and COVID19 patients (C) Relative mRNA expression levels of inflammatory biomarkers and transcription factors in the lung biopsies of healthy and COVID19 patients. Two independent samples for each condition were shown.

**Figure 14.**
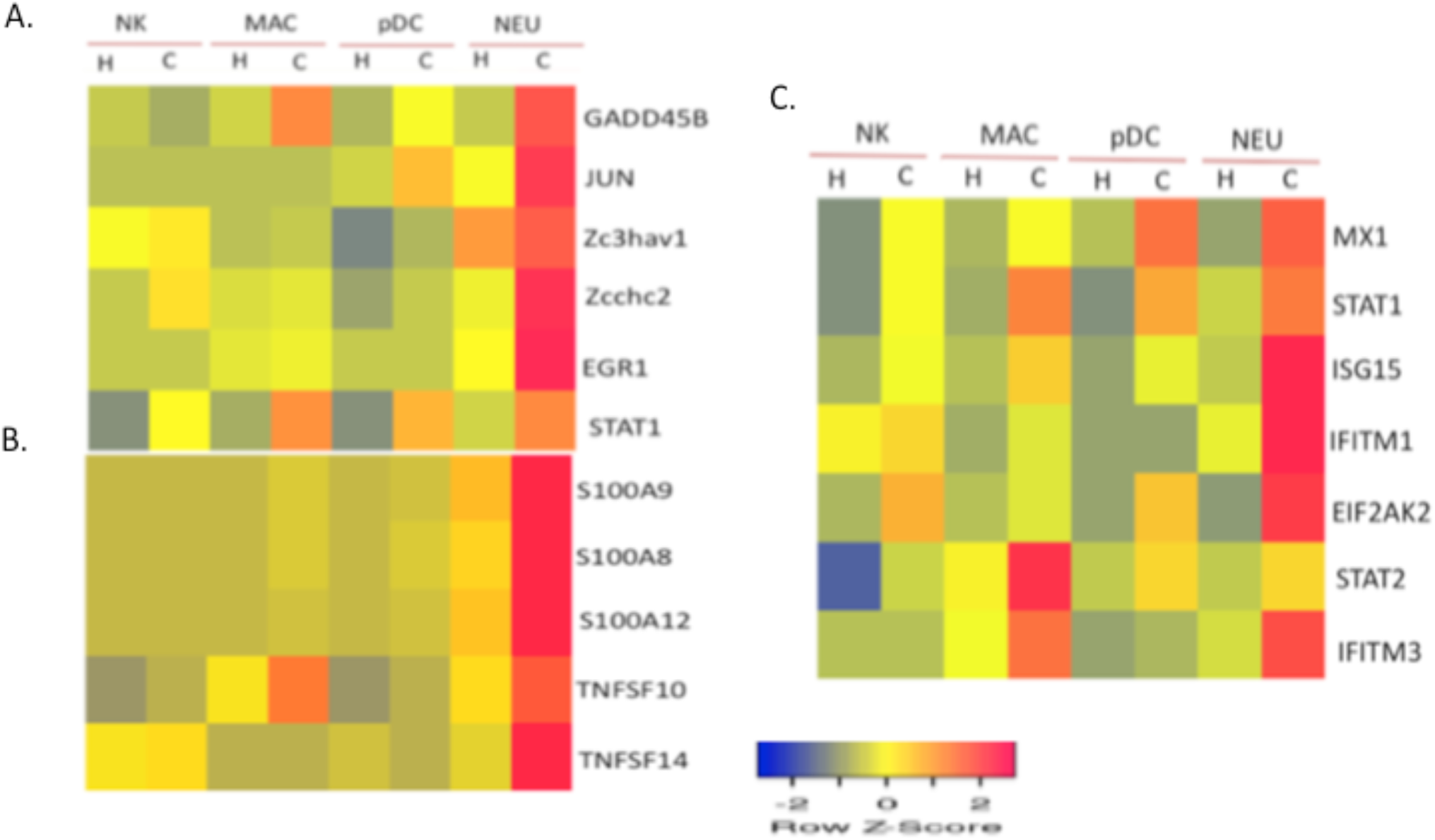
Regulation of transcription factor and inflammatory biomarker mRNA levels by type I interferon signaling in the PBMC of healthy and COVID19 patients (A) Expression levels of transcription factors of the antiviral, immune modulation, and cell growth sub-networks in the PBMC of healthy and COVID19 patients (B) Expression levels of inflammatory biomarkers in the PBMC of healthy and COVID19 patients (C) Cluster analysis of transcription factors and effectors of antiviral sub-network in the PBMC of healthy and COVID19 patients. Natural killer cells (NK), macrophages (MAC, CD16 positive), dendritic cells (pDC), and neutrophils (Neu) in Healthy (H) and COVID19 patients (C) were shown.

## 4 Conclusion

Genetic variants or deficiency in components of type I interferon signaling was reported in a significant proportion of intensive care COVID-19 patients (21-23). In contrast, enhanced inflammatory response mediated by interferons and cytokines was also reported ln COVID-19 patients leading to an unbalanced cytokine response (27,70-72). Transcriptional factor profiling revealed that there are two distinct steps in the transcriptional regulation by type I interferons-a fast-acting tyrosine phosphorylation switch leading to the activation of the ISGF-3 complex and a gradual increase of inducible transcription factors that regulate the secondary and tertiary transcription programs. These two pathways overlap and co-regulate distinct sets of genes to confer temporal regulation of the type I interferon response. Distinct sub-networks of transcription factors including antiviral response, immune modulation, and cell growth mediate the biological effects in type I interferon signaling. Differential regulation of sub-networks was observed in human lung epithelial cells after SARS-CoV-2 infection and in COVID-19 patients. Furthermore, a gene signature of type I interferon signaling transcription factors and inflammatory mediators was observed in the lungs and peripheral blood of COVID-19 patients. The topology of the interferon regulated transcription factor sub-networks and their target gene expression may be critically assessed using software tools developed for social network analysis. Type I interferons have pleiotropic effects and are promising therapeutic agents in the treatment of different classes of diseases such as viral infection, autoimmune disease, and cancer. A detailed understanding of the IFN α/β signaling mechanisms is important for the effectiveness of the therapies. Transcription factor profiling in type I interferon signaling may have therapeutic implications for treating COVID-19 patients by targeting selective inflammatory pathways such as MAP kinase pathway and Early growth response 1 (Egr-1) involved in the regulation of chemokines and cell growth (26). In addition, transcription factor profiling of target genes involved in innate and adaptive immunity revealed that critical nodes such as BRCA1 and GILZ involved in regulating multiple cytokine signaling pathways can be simultaneously targeted with small molecules such as dexamethasone to attenuate inflammation in lung airway epithelial cells (25). Furthermore, COVID-19 patients with high interferon-stimulated gene expression (ISG^high^) in the lungs die significantly earlier than patients with low ISG expression (ISG^low^) after hospitalization (72). In this report, high-levels of expression of a subset of type I interferon-inducible transcription factors and genes involved in cell apoptosis and inflammation were observed in the lungs and neutrophils in COVID-19 patients. The use of Jak1 or Tyk2 selective inhibitors to block type I interferon signaling in the periphery and lungs or locally to lower ISG expression may have therapeutic applications in the treatment of critically ill COVID-19 patients (81,82).

## Acknowledgements

I would like to thank Dr. Donald Forsdyke (Queen’s University, Canada) for a careful reading of the manuscript and N.S. Sastry (Chelmsford, MA, USA) for continuous support.

## REFERENCES

1. Stark GR, Kerr IM, Williams BR et al. How cells respond to interferons. Annu Rev Biochem. 1998; 67:227–264.

2. Park A, Iwasaki A. Type I and Type III Interferons - Induction, Signaling, Evasion, and Application to Combat COVID-19. Cell Host Microbe. 2020; 27:870–878.

3. Platanias L Mechanisms of type-I- and Type-II-interferon-mediated signaling. Nature Rev Immunol. 2005; 5:375–386.

4. Weber F, Haller O. Viral suppression of the interferon system. Biochimie. 2007; 89:836–42.

5. Der SD, Zhou A, Williams BR, Silverman RH. Identification of genes differentially regulated by interferon alpha, beta, or gamma using oligonucleotide arrays. Proc Natl Acad Sci U S A. 1998; 95:15623–15628.

6. Stark GR, Darnell JE Jr. The JAK-STAT pathway at twenty. Immunity. 2012; 36:503–514.

7. Waddell SJ, Popper SJ, Rubins KH, et al. Dissecting interferon-induced transcriptional programs in human peripheral blood cells. PLoS One. 2010; 5:e9753.

8. Mostafavi S, Yoshida H, Moodley D, LeBoité H, Rothamel K, Raj T, Ye CJ, Chevrier N, Zhang SY, Feng T, Lee M, Casanova JL, Clark JD, Hegen M, Telliez JB, Hacohen N, De Jager PL, Regev A, Mathis D, Benoist C; Immunological Genome Project Consortium. Parsing the Interferon Transcriptional Network and Its Disease Associations. Cell. 2016; 164:564–78.

9. Gongora R, Stephan RP, Schreiber RD, Cooper MD. Stat-1 is not essential for inhibition of B lymphopoiesis by type I IFNs. J Immunol. 2000; 165:2362–2366.

10. Gongora R, Stephan RP, Zhang Z, Cooper MD. An essential role for Daxx in the inhibition of B lymphopoiesis by type I interferons. Immunity. 2001;14:727–737.

11. Meraz MA, White JM, Sheehan KC, Bach EA, Rodig SJ, Dighe AS, Kaplan DH, Riley JK, Greenlund AC, Campbell D, Carver-Moore K, DuBois RN, Clark R, Aguet M, Schreiber RD. Targeted disruption of the Stat1 gene in mice reveals unexpected physiologic specificity in the JAK-STAT signaling pathway. Cell. 1996; 84:431–442.

12. Durbin JE, Hackenmiller R, Simon MC, Levy DE. Targeted disruption of the mouse Stat1 gene results in compromised innate immunity to viral disease. Cell. 1996; 84:443–450.

13. Gil MP, Bohn E, O’Guin AK, Ramana CV, Levine B, Stark GR, Virgin HW, Schreiber RD. Biologic consequences of Stat1-independent IFN signaling. Proc Natl Acad Sci U S A. 2001; 98:6680–6685.

14. Wang J, Schreiber RD, Campbell IL. STAT1 deficiency unexpectedly and markedly exacerbates the pathophysiological actions of IFN-alpha in the central nervous system. Proc Natl Acad Sci U S A. 2002; 99:16209–16214.

15. Ousman SS, Wang J, Campbell IL. Differential regulation of interferon regulatory factor (IRF)-7 and IRF-9 gene expression in the central nervous system during viral infection. J Virol. 2005; 79:7514–7527.

16. Kohlmeier JE, Woodland DL. Immunity to respiratory viruses. Annu Rev Immunol. 2009; 27:61–82.

17. Ardain A, Marakalala MJ, Leslie A. Tissue-resident innate immunity in the lung. Immunology. 2020; 159:245–256.

18. Cui J, Li F, Shi ZL. Origin and evolution of pathogenic coronaviruses. Nat Rev Microbiol. 2019;17(3):181–192.

19. Zhou, P., Yang, X.-L., Wang, X.-G., Hu, B., Zhang, L et. al. (2020). A pneumonia outbreak associated with a new coronavirus of probable bat origin. Nature. 579: 270–273.

20. Huang, C., Wang, Y., Li, X., et.al. (2020). Clinical features of patients infected with 2019 novel coronavirus in Wuhan, China. The Lancet. 395:497–506.

21. Zhang Q, Bastard P, Liu Z, Le Pen J, Moncada-Velez M, Chen J, et.al. Inborn errors of type I IFN immunity in patients with life-threatening COVID-19. Science. 2020; 370:eabd4570

22. Bastard P, Rosen LB, Zhang Q, Michailidis E, Hoffmann HH, et. al. Autoantibodies against type I IFNs in patients with life-threatening COVID-19. Science. 2020; 370(6515)

23. Pairo-Castineira E, Clohisey S, Klaric L, Bretherick AD, Rawlik K,. Genetic mechanisms of critical illness in Covid-19. Nature. 2020. doi: 10.1038/s41586-020-03065-y.

24. Ramana CV. Insights into the signal transduction pathways of mouse lung type II cells revealed by Transcription Factor Profiling in the transcriptome. Genomics Inform. 2019; 17(1):e8.

25. Ramana CV. Insights into functional connectivity in mammalian signal transduction pathways by pairwise comparison of protein interaction partners of critical signaling hubs. bioRxiv 2019. DOI: 10.1101/2019.12.30.891200

26. Ramana C.V. Regulation of early growth response-1 (Egr-1) gene expression by Stat1-independent type I interferon signaling and respiratory viruses. bioRxiv 2020. DOI: 10.1101/2020.08.14.244897

27. Blanco-Melo D, Nilsson-Payant BE, Liu WC, Uhl S, Hoagland D, Møller R, Jordan TX, Oishi K, Panis M, Sachs D, Wang TT, Schwartz RE, Lim JK, Albrecht RA, tenOever BR. Imbalanced Host Response to SARS-CoV-2 Drives Development of COVID-19. Cell. 2020; 181:1036-1045.e9.

28. Wilk AJ, Rustagi A, Zhao NQ, Roque J, Martínez-Colón GJ, McKechnie JL, Ivison GT, Ranganath T, Vergara R, Hollis T, Simpson LJ, Grant P, Subramanian A, Rogers AJ, Blish CA. A single-cell atlas of the peripheral immune response in patients with severe COVID-19. Nat Med. 2020; 7:1070–1076.

29. Babicki S, Arndt D, Marcu A, Liang Y, Grant JR, Maciejewski A, Wishart DS. Heatmapper: web-enabled heat mapping for all. Nucleic Acids Res. 2016; 44(W1):W147–53.

30. Huang da W, Sherman BT, Lempicki RA. Systematic and integrative analysis of large gene lists using DAVID bioinformatics resources. Nat Protoc. 2009; 4(1):44.

31. Kanehisa M, Furumichi M, Tanabe M, Sato Y, Morishima K. KEGG: new perspectives on genomes, pathways, diseases and drugs. Nucleic Acids Res. 2017; 45:D353–D361.

32. Zhou Y, Zhou B, Pache L, Chang M, Khodabakhshi AH, Tanaseichuk O, Benner C, Chanda SK. Metascape provides a biologist-oriented resource for the analysis of systems-level datasets. Nat Commun. 2019; 10:1523

33. Stark C, Breitkreutz BJ, Reguly T, Boucher L, Breitkreutz A, Tyers M. BIOGRID: a general repository for interaction datasets. Nucleic Acids Res. 2006; 34:D535–D539.

34. Szklarczyk D, Morris JH, Cook H, Kuhn M, Wyder S, Simonovic M, Santos A, Doncheva NT, Roth A, Bork P, Jensen LJ, von Mering C. The STRING database in 2017. Quality-controlled protein-protein association networks, made broadly accessible. Nucleic Acids Res. 2017; 45:D362–D368.

35. Orii N, Ganapathiraju MK. WIKI-PI: a web-server of annotated human protein-protein interactions to aid in discovery of protein function PLoS One. 2012; 7:e49029.

36. Basler CF, García-Sastre A. Viruses and the type I interferon antiviral system: induction and evasion. Int Rev Immunol. 2002; 2:305–337.

37. Dupuis S, Dargemont C, Fieschi C, Thomassin N, Rosenzweig S, Harris J, Holland SM, Schreiber RD, Casanova JL. Impairment of mycobacterial but not viral immunity by a germline human STAT1 mutation. Science. 2001; 293:300–303

38. Mavrommatis E, Fish EN, Platanias LC. The schlafen family of proteins and their regulation by interferons. J Interferon Cytokine Res. 2013; 33:206–210.

39. Andersson R, Sandelin A. Determinants of enhancer and promoter activities of regulatory elements. Nat Rev Genet. 2020; 21:71–87.

40. Papin JA, Hunter T, Palsson BO, Subramaniam S., Reconstruction of cellular signaling networks and analysis of their properties. Nat Rev Mol Cell Biol. 2005; 6:99–111.

41. Korzus E, Torchia J, Rose DW, Xu L, Kurokawa R, McInerney EM, Mullen TM, Glass CK, Rosenfeld MG. Transcription factor-specific requirements for coactivators and their acetyltransferase functions. Science. 1998; 279:703–707.

42. Zhang JJ, Vinkemeier U, Gu W, Chakravarti D, Horvath CM, Darnell JE Jr. Two contact regions between Stat1 and CBP/p300 in interferon gamma signaling. Proc Natl Acad Sci U S A. 1996; 93:15092–15096.

43. Zeng L, Sun Y, Xie L, Wei L, Ren Y, Zhao J, Qin W, Mitchelson K, Cheng J. Construction of a novel oligonucleotide array-based transcription factor interaction assay platform and its uses for profiling STAT1 cofactors in mouse fibroblast cells. Proteomics. 2013;13:2377–2385.

44. Bulgakov VP, Tsitsishvili GSh. Bioinformatics analysis of protein interaction networks: statistics, topologies, and meeting the standards of experimental biologists. Biochemistry (Mosc). 2013; 78:1098–103.

45. Han H, Cho JW, Lee S, Yun A, Kim H, Bae D, Yang S, Kim CY, Lee M, Kim E, Lee S, Kang B, Jeong D, Kim Y, Jeon HN, Jung H, Nam S, Chung M, Kim JH, Lee I. TRRUST v2: an expanded reference database of human and mouse transcriptional regulatory interactions Nucleic Acids Res. 2018; 46:D380–D386.

46. Ashtiani M, Salehzadeh-Yazdi A, Razaghi-Moghadam Z, Hennig H, Wolkenhauer O, Mirzaie M, Jafari M. A systematic survey of centrality measures for protein-protein interaction networks. BMC Syst Biol. 2018; 12:80.

47. Kleinberg, J. Authoritative sources in a hyper-linked environment. Journal of the ACM. 1999; 46:604–632.

48. Adelaja A, Hoffmann A. Signaling Crosstalk Mechanisms That May Fine-Tune Pathogen-Responsive NFκB. Front Immunol. 2019; 10:433.

49. Langevin C, Aleksejeva E, Passoni G, Palha N, Levraud JP, Boudinot P. The antiviral innate immune response in fish: evolution and conservation of the IFN system. J Mol Biol. 2013; 425:4904–4920.

50. Majzoub K, Wrensch F, Baumert TF. The Innate Antiviral Response in Animals: An Evolutionary Perspective from Flagellates to Humans. Viruses. 2019; 11:758.

51. Shaw AE, Hughes J, Gu Q, Behdenna A, Singer JB, Dennis T, Orton RJ, Varela M, Gifford RJ, Wilson SJ, Palmarini M. Fundamental properties of the mammalian innate immune system revealed by multispecies comparison of type I interferon responses. PLoS Biol. 2017;15:e2004086

52. Zhang B, Goraya MU, Chen N, Xu L, Hong Y, Zhu M, Chen JL. Zinc Finger CCCH-Type Antiviral Protein 1 Restricts the Viral Replication by Positively Regulating Type I Interferon Response. Front Microbiol. 2020; 11:1912.

53. Dai H, Yan M, Li Y. The zinc-finger protein ZCCHC2 suppresses retinoblastoma tumorigenesis by inhibiting HectH9-mediated K63-linked polyubiquitination and activation of c-Myc. Biochem Biophys Res Commun. 2020; 521:533–538.

54. Ramana CV, Chatterjee-Kishore M. Nguyen H, Stark GR. Complex roles of Stat1 in regulating gene expression. Oncogene. 2000; 19:2619–2627.

55. Velichko S, Wagner TC, Turkson J, Jove R, Croze E. STAT3 activation by type I interferons is dependent on specific tyrosines located in the cytoplasmic domain of interferon receptor chain 2c. Activation of multiple STATS proceeds through the redundant usage of two tyrosine residues. J Biol Chem. 2002;277:35635–35641.

56. Watling D, Guschin D, Müller M, Silvennoinen O, Witthuhn BA, Quelle FW, Rogers NC, Schindler C, Stark GR, Ihle JN, et al. Complementation by the protein tyrosine kinase JAK2 of a mutant cell line defective in the interferon-gamma signal transduction pathway. Nature. 1993; 366:166–170.

57. Karaghiosoff M, Neubauer H, Lassnig C, Kovarik P, Schindler H, Pircher H, McCoy B, Bogdan C, Decker T, Brem G, Pfeffer K, Müller M. Partial impairment of cytokine responses in Tyk2-deficient mice. Immunity. 2000 ;13:549–560.

58. Yang CH, Murti A, Pfeffer LM. STAT3 complements defects in an interferon-resistant cell line: evidence for an essential role for STAT3 in interferon signaling and biological activities. Proc Natl Acad Sci U S A. 1998; 95:5568–72.

59. Farrar JD, Smith JD, Murphy TL, Leung S, Stark GR, Murphy KM. Selective loss of type I interferon-induced STAT4 activation caused by a minisatellite insertion in mouse Stat2. Nat Immunol. 2000;1:65–69.

60. Gadina M, O’Shea JJ, Biron CA. Critical role for STAT4 activation by type 1 interferons in the interferon-gamma response to viral infection. Science. 2002; 297:2063–2066.

61. Nguyen KB, Salazar-Mather TP, Dalod MY, Van Deusen JB, Wei XQ, Liew FY, Caligiuri MA, Durbin JE, Biron CA. Coordinated and distinct roles for IFN-alpha beta, IL-12, and IL-15 regulation of NK cell responses to viral infection. J Immunol. 2002; 169:4279–4287

62. Matikainen S, Sareneva T, Ronni T, Lehtonen A, Koskinen PJ, Julkunen I. Interferon-alpha activates multiple STAT proteins and upregulates proliferation-associated IL-2Ralpha, c-myc, and pim-1 genes in human T cells. Blood. 1999; 93:1980–1991.

63. Meinke A, Barahmand-Pour F, Wöhrl S, Stoiber D, Decker T. Activation of different Stat5 isoforms contributes to cell-type-restricted signaling in response to interferons. Mol Cell Biol. 1996; 16:6937–6944.

64. Rani MR, Leaman DW, Han Y, Leung S, Croze E, Fish EN, Wolfman A, Ransohoff RM. Catalytically active TYK2 is essential for interferon-beta-mediated phosphorylation of STAT3 and interferon-alpha receptor-1 (IFNAR-1) but not for activation of phosphoinositide 3-kinase. J Biol Chem. 1999; 274:32507–32511.

65. Dupuis S, Jouanguy E, Al-Hajjar S, Fieschi C, Al-Mohsen IZ, Al-Jumaah S, Yang K, Chapgier A, Eidenschenk C, Eid P, Al Ghunaim A, Tufenkji H, Frayha H, Al-Gazlan S, Al-Rayes H, Schreiber RD, Gresser I, Casanova JL. Impaired response to interferon-alpha/beta and lethal viral disease in human STAT1 deficiency. Nat Genet. 2003; 33:388–391.

66. Tsai MH, Pai LM, Lee CK. Fine-Tuning of Type I Interferon Response by STAT3. Front Immunol. 2019; 10:1448

67. Uddin S, Lekmine F, Sassano A, Rui H, Fish EN, Platanias LC. Role of Stat5 in type I interferon-signaling and transcriptional regulation. Biochem Biophys Res Commun. 2003; 308:325–330.

68. Pfeffer LM, Mullersman JE, Pfeffer SR, Murti A, Shi W, Yang CH. STAT3 as an adapter to couple phosphatidylinositol 3-kinase to the IFNAR1 chain of the type I interferon receptor. Science. 1997; 276:1418–1420.

69. Wagner TC, Velichko S, Vogel D, Rani MR, Leung S, Ransohoff RM, Stark GR, Perez HD, Croze E. Interferon signaling is dependent on specific tyrosines located within the intracellular domain of IFNAR2c. Expression of IFNAR2c tyrosine mutants in U5A cells. J Biol Chem. 2002; 277:1493–1499.

70. Arunachalam PS, Wimmers F, Mok CKP, Perera RAPM, Scott M, Hagan T, Sigal N, Feng Y, Bristow L, Tak-Yin Tsang O, Wagh D, Coller J, Pellegrini KL, Kazmin D, Alaaeddine G, Leung WS, Chan JMC, Chik TSH, Choi CYC, Huerta C, Paine McCullough M, Lv H, Anderson E, Edupuganti S, Upadhyay AA, Bosinger SE, Maecker HT, Khatri P, Rouphael N, Peiris M, Pulendran B. Systems biological assessment of immunity to mild versus severe COVID-19 infection in humans. Science. 2020; 369:1210–1220.

71. Lucas C, Wong P, Klein J, Castro TBR, Silva J, Sundaram M, Ellingson MK, Mao T, Oh JE, Israelow B, Takahashi T, Tokuyama M, Lu P, Venkataraman A, Park A, Mohanty S, Wang H, Wyllie AL, Vogels CBF, Earnest R, Lapidus S, Ott IM, Moore AJ, Muenker MC, Fournier JB, Campbell M, Odio CD, Casanovas-Massana A; Yale IMPACT Team, Herbst R, Shaw AC, Medzhitov R, Schulz WL, Grubaugh ND, Dela Cruz C, Farhadian S, Ko AI, Omer SB, Iwasaki A. Longitudinal analyses reveal immunological misfiring in severe COVID-19. Nature. 2020; 584:463–469.

72. Nienhold R, Ciani Y, Koelzer VH, Tzankov A, Haslbauer JD, Menter T, Schwab N, Henkel M, Frank A, Zsikla V, Willi N, Kempf W, Hoyler T, Barbareschi M, Moch H, Tolnay M, Cathomas G, Demichelis F, Junt T, Mertz KD. Two distinct immunopathological profiles in autopsy lungs of COVID-19. Nat Commun. 2020;11:5086.

73. Xu L, Yoon H, Zhao MQ, Liu J, Ramana CV, Enelow RI. Cutting edge: pulmonary immunopathology mediated by antigen-specific expression of TNF-alpha by antiviral CD8+ T cells. J Immunol. 2004; 173:721–725.

74. Ramana, CV, DeBerge MP, Kumar A, Alia CS, Durbin JE, Enelow RI. Inflammatory impact of IFN-γ in CD8+ T cell-mediated lung injury is mediated by both Stat1-dependent and – independent pathways. Am J Physiol Lung Cell Mol Physiol. 2015; 308:L650–L657.

75. Verdecchia P, Cavallini C, Spanevello A, Angeli F. The pivotal link between ACE2 deficiency and SARS-CoV-2 infection. Eur J Intern Med. 2020; 76:14–20.

76. Imgenberg-Kreuz J, Sandling JK, Björk A, Nordlund J, Kvarnström M, Eloranta ML, Rönnblom L, Wahren-Herlenius M, Syvänen AC, Nordmark G. Transcription profiling of peripheral B cells in antibody-positive primary Sjögren’s syndrome reveals upregulated expression of CX3CR1 and a type I and type II interferon signature. Scand J Immunol. 2018; 87:e12662.

77. Imgenberg-Kreuz J, Sandling JK, Almlöf JC, Nordlund J, Signér L, Norheim KB, Omdal R, Rönnblom L, Eloranta ML, Syvänen AC, Nordmark G. Genome-wide DNA methylation analysis in multiple tissues in primary Sjögren’s syndrome reveals regulatory effects at interferon-induced genes. Ann Rheum Dis. 2016; 75: 2029–2036.

78. Silvin A, Chapuis N, Dunsmore G, Goubet AG, Dubuisson A, Derosa L, Almire C, Hénon C, Kosmider O, Droin N, Rameau P, Catelain C, Alfaro A, Dussiau C, Friedrich C, Sourdeau E, Marin N, Szwebel TA, Cantin D, Mouthon L, Borderie D, Deloger M, Bredel D, Mouraud S, Drubay D, Andrieu M, L’honneur AS, Saada V, Stoclin A, Willekens C, Pommeret F, Griscelli F, Ng LG, Zhang Z, Bost P, Amit I, Barlesi F, Marabelle A, Pène F, Gachot B, André F, Zitvogel L, Ginhoux F, Fontenay M, Solary E. Elevated Calprotectin and Abnormal Myeloid Cell Subsets Discriminate Severe from Mild COVID-19. Cell. 2020; 182:1401-1418.e18.

79. Croft M, Duan W, Choi H, Eun SY, Madireddi S, Mehta A. TNF superfamily in inflammatory disease: translating basic insights. Trends Immunol. 2012; 33:144–52.

80. Xia C, Braunstein Z, Toomey AC, Zhong J, Rao X. S100 Proteins as an important regulator of macrophage inflammation. Front Immunol. 2018; 8:1908.

81. Wrobleski ST, Moslin R, Lin S, Zhang Y, Spergel S, Kempson J, Tokarski JS, Strnad J, Zupa-Fernandez A, Cheng L, Shuster D, Gillooly K, Yang X, Heimrich E, McIntyre KW, Chaudhry C, Khan J, Ruzanov M, Tredup J, Mulligan D, Xie D, Sun H, Huang C, D’Arienzo C, Aranibar N, Chiney M, Chimalakonda A, Pitts WJ, Lombardo L, Carter PH, Burke JR, Weinstein DS. Highly Selective Inhibition of Tyrosine Kinase 2 (TYK2) for the Treatment of Autoimmune Diseases: Discovery of the Allosteric Inhibitor BMS-986165. J Med Chem. 2019 Oct 24;62(20):8973–8995.

82. Norman P. Selective JAK1 inhibitor and selective Tyk2 inhibitor patents. Expert Opin Ther Pat. 2012;22:1233–1249.

